# CTCF-DEPENDENT INSULATION OF *Hoxb13* AND THE HETEROCHRONIC CONTROL OF TAIL LENGTH

**DOI:** 10.1101/2024.07.20.604409

**Authors:** Lucille Lopez-Delisle, Jozsef Zakany, Célia Bochaton, Pierre Osteil, Alexandre Mayran, Fabrice Darbellay, Bénédicte Mascrez, Hocine Rekaik, Denis Duboule

## Abstract

In mammals, tail length is controlled by several genetic determinants, amongst which Hox13 genes located at the posterior extremities of Hox clusters, whose main function are to terminate the extension of the body axis. In this view, the precise timing in the transcriptional activation of these genes may impact upon body length. Unlike other Hox clusters, HoxB lacks all posterior genes between Hoxb9 and Hoxb13, two genes separated by a ca. 70 kb large DNA segment containing an unusually high number of CTCF sites, suggesting it isolates Hoxb13 from the rest of the cluster, thereby delaying its negative impact on trunk extension. We deleted the spacer DNA to induce a potential heterochronic gain of function of Hoxb13 at physiological concentration and observed a shortening of the tail as well as other abnormal phenotypes, which were all rescued by inactivating Hoxb13 in-cis with the deletion. A comparable gain of function was observed in mutant ES cells grown as pseudo-embryos in vitro, which allowed us to examine in details the importance of both the number and the orientation of CTCF sites in the insulating activity of the DNA spacer. A short cassette containing all the CTCF sites was sufficient to insulate Hoxb13 from the rest of HoxB and additional modifications of this CTCF cassette showed that two CTCF sites in convergent orientations are already capable of importantly delaying Hoxb13 activation in these conditions. We discuss the relative importance of genomic distance versus number and orientation of CTCF sites in preventing Hoxb13 to be activated too early during trunk extension and hence to modulate tail length.

## INTRODUCTION

The body axis of most mammalian species including humans (1) terminates with a tail of a defined and characteristic length. Since the isolation of the T/Brachyury gene (2), several genetic determinants of tail variation have been described and progresses have been made in exploring the mechanisms of length variability either during development (3), in wild mouse populations (4) or during human evolution (5). In this context, several studies reported a role for the most posterior *Hox13* genes in setting up tail length (4, 6) and a genetic modification in mice (7) produced tail overgrowth reminiscent of the targeted inactivation of *Hoxb13* (8, 9). Conversely, forced expression of either *Hoxb13* or of other group 13 genes induced variable, often dramatic vertebral column truncations (10). As a consequence, it was proposed that *Hox13* genes were collectively responsible for terminating the extension of the main body axis during embryogenesis (10–12), thereby setting the length of the tail through a dominant negative effect of their protein products (13). Recent results obtained using unbiased approaches in natural populations of either deer mice (4), or Chinese long-tailed sheep breeds (14) have further identified *Hoxd13* and *Hoxb13* as candidate genes to regulate tail length.

Since *HOX13* proteins participate to the termination of the major body axis, it is important that these genes are kept silent until the time and the place when they need to be implemented such as to avoid premature termination. This is at least partly controlled by a tight mechanism that involves the separation of group 13 genes from the rest of the *Hox* clusters due to the presence of a chromatin border between two topologically associating domains (TADs) (15, 16), which insulate *Hox13* genes from regulatory influences emanating from the neighbouring TAD (17, 18). Such TAD borders are often organized by the presence of several CTCF binding sites with opposite orientations leading to separate tropisms for chromatin loop extrusion (see (19)). This somewhat generic organisation is particularly visible and conserved within the *HoxA*, *HoxC* and *HoxD* gene clusters (18, 20), which likely reflects an ancestral chromatin architecture present before the occurrence of the full genomic duplications that generated these multiple clusters.

The *HoxB* cluster, on the other hand, shows a somewhat distinct topology since it lacks the *Hoxb10*, *Hoxb11* and *Hoxb12* genes, i.e., a region that contains several such CTCF sites in the other three clusters. However, *Hoxb13* is not found at the usually close vicinity of its nearest neighbour *Hoxb9*, but instead it lies ca. 70 kb far from it (8), which explains why it was initially overlooked when cloning the *HoxB* clusters in human and mice (21, 22). Furthermore, several CTCF binding sites (CBSs) are found regularly spaced within this DNA segment (Fig. 1A, CBS5 to CBS10), as if *Hox* genes had been deleted after genome duplications while leaving in place their associated CBSs (17, 18). This suggests that the length of the spacer DNA segment and/or its content in CBSs participate in the insulation of *Hoxb13* and thus delays its timing of activation. This hypothesis is supported by chromosome conformation capture experiments showing that this DNA spacer acts as a chromatin boundary between the *Hoxb1* to *Hoxb9* region, on the one hand, and *Hoxb13* on the other hand (23), as expected from the orientations of five out of six CTCF sites present in the DNA spacer (18).

**Figure 1.**
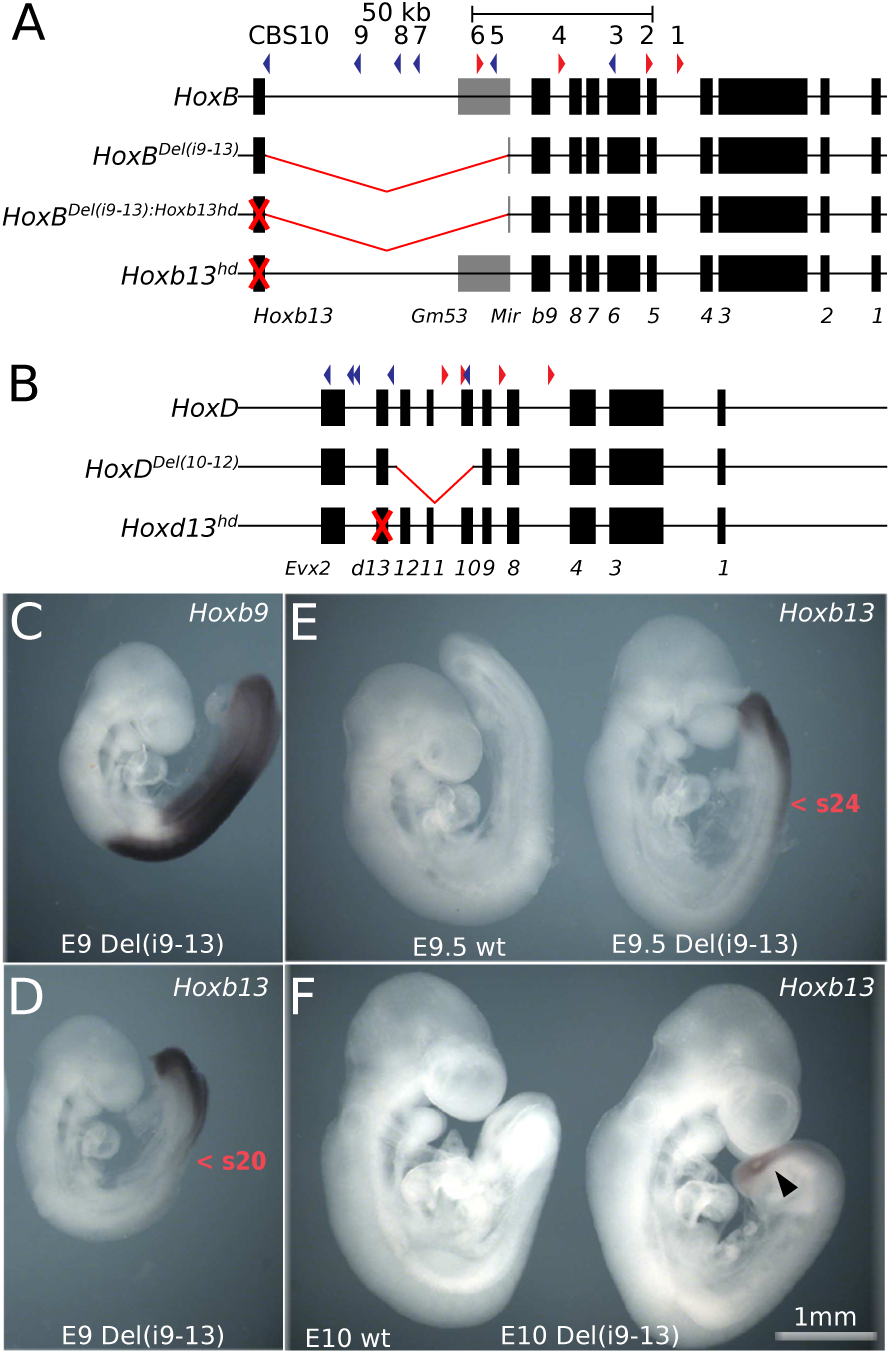
Deletion of the *Hoxb9* to *Hoxb13* spacer DNA in the *HoxB* cluster. (**A**) Schematic representation of the *HoxB* alleles used in this work. On top, the CTCF binding sites (CBS) are numbered 1 to 10 from *Hoxb1* to *Hoxb13* and the coloured arrowheads below indicate orientations. Below, the wild type *HoxB* locus is shown. An approximately 70 kb large region separates *Hoxb9* from *Hoxb13*, without any protein coding genes, whereas a LncRNA (*Gm53*) and the microRNA *Mir196a-1* (’Mir’) are found within the 19 kb immediately flanking *Hoxb9*. The size of this spacer DNA is well conserved in all investigated species (see Fig. S1). As in all amniotes, *Hoxb10*, *11* and *12* are absent. The extent of the induced *HoxB(i9-13)* deletion is shown below, as well as the loss of function *Hoxb13^hd^* allele generated either *in-cis* with the *HoxB(i9-13)* deletion, or *in-trans* (red crosses). (**B**) Schematic representation of the *HoxD* alleles used in this work (same scale as *HoxB*). On top the wild type *HoxD* locus is shown. The extent of the *Del*(*10–12*) deletion bringing *Hoxd13* in close proximity to *Hoxd9* (25) is shown below, as well as the loss of function *Hoxd13^hd^* allele (red cross) (26). (**C**) WISH using a *Hoxb9* probe at E9. *Hoxb9* expression pattern was indistinguishable between wild type and homozygous *HoxB^Del(i9-13)^* mutant specimens. (**D**) *Hoxb13* transcript accumulation at E9 in a homozygous *HoxB^Del(i9-13)^* mutant specimen. At this stage *Hoxb13* signal was undetectable in wild type littermates. The red arrowhead highlights the position of somite 20. (**E**) *Hoxb13* transcript accumulation in a E9.5 homozygous *HoxB^Del(i9-13)^* mutant specimen (right), compared with a control littermate (left) still negative for *Hoxb13* mRNAs. The red arrowhead highlights the position of somite 24. (**F**) *Hoxb13* expression in a homozygous *HoxB^Del(i9-13)^* mutant specimen at E10 (right) compared to a control littermate (left). The expression of *Hoxb13* is highlighted by the black arrowhead. All specimens were treated in the same experiment in parallel, using the same stocks of reagents, incubations and washes. The arrowhead shows expression in the mutant specimen only.

To verify the importance of this region in delaying *Hoxb13* expression in the elongating trunk, we shortened it from ca. 70 kb down to 6.6 kb such as to bring *Hoxb13* near *Hoxb9* by removing at the same time all CBSs present in this ‘spacer’ DNA. We show that this condensed *HoxB* cluster leads to mice with short tails assorted with additional thoracic and lumbar vertebra losses. A secondary targeted inactivation of *Hoxb13 in-cis* rescues all vertebral column defects demonstrating that the gained HOXB13 protein is solely responsible for the abnormal phenotypes, while transcriptome studies suggest that tail shortening is due to the precocious activation of a *Hoxb13* functional program. To discriminate between the importance of the length of the DNA spacer *versus* the presence of multiple CBSs in the insulation of *Hoxb13*, we reproduced the same deletion in ES-cells derived pseudo-embryos. Premature *Hoxb13* activation was observed and a synthetic cassette containing all or some CTCF sites was recombined between *Hoxb9* and *Hoxb13* to see its impact upon keeping *Hoxb13* silent.

## RESULTS

We induced a 67.5 kb large deletion of the mouse *HoxB* intergenic spacer region by using CRISPR/Cas9 (the *HoxB^Del(i9-13)^* allele, Fig. 1A). At the breakpoint near the *HoxB* cluster, the deletion included the *GM53* LncRNA of unknown function, yet the *Mir196a-1* was left in place due to its potential importance in regulating some *Hox* RNAs stability ((24) and ref. therein). A survey of 157 informative genome sequences revealed that this short distance between *Hoxb13* and *Hoxb9* (9.3 kb) is two times smaller than that observed in some rare fish genomes, while the shortest distances found in amniotes is four times this large in aves and at least six times as large in placentalia (Fig. S1A). Also, while the length of this spacer is globally maintained throughout mammals, the DNA sequence is poorly conserved (Fig. S1B), even when mouse inbred strains are compared (Fig. S1C), suggesting that a minimal DNA length between *Hoxb9* and *Hoxb13* had been selected. We also produced two control alleles where we inactivated *Hoxb13* function either *in-cis* with the spacer deletion (*HoxB^Del(i9-13):Hoxb13hd^*), or *in-trans* (*Hoxb13^hd^*, Fig. 1A). The related *HoxD^Del^*(*^10–12^*) and *Hoxd13^hd^* alleles (25, 26) were also investigated in this context to test for a potential synergy in phenotypes (Fig. 1B).

### Heterochronic shift of *Hoxb13* expression in the deletion mutant

Whole mount *in situ* hybridization (WISH) on *HoxB^Del(i9-13)^* mutant embryos showed strong *Hoxb9* expression at E9 (Theiler stage 14, Fig. 1C), identical to the wt expression, whereas *Hoxb13* transcripts were not yet detected (Fig. 1E, left). In contrast, expression of *Hoxb13* was detected as early as E9 in *HoxB^Del(i9-13)^* mutant embryos (Fig. 1D). The signal was localized to the posterior trunk, with an anterior limit approximately at somite 20 (Fig. 1, red arrowhead), which contributes to the formation of pre-vertebra 16 i.e., thoracic vertebra 9 (T9). Later, the anterior limit of *Hoxb13* signal had shifted to ca. somite number 24 (Fig. 1E, right). In formed epithelial somites, signal was weak, if any, compared to the posterior pole of the embryo including the pre-somitic mesoderm (PSM). In late E10 embryos, a weak ectopic *Hoxb13* signal was still visible in the tail around the posterior neuropore (Fig. 1F, right, arrowhead). All these *Hoxb13* signals were clearly premature and ectopic. Indeed, the wild type *Hoxb13* signal was first detected at somite level 45 at E11, Theiler stage 18, which corresponds to pre-vertebra 41 i.e., the eleventh caudal vertebra (CA11, not shown). These results were confirmed by RNAseq analysis (see below).

### Vertebral column malformations in adult *HoxB^Del(i9-13)^* mutant mice

A gross observation of *HoxB^Del(i9-13)^*mutant specimen (or Del(i9-13)) revealed a tail shorter than in wild type littermates and measuring the distance between the anus and the tail tip in F2 adults quantified these distinct tail truncations, an observation controlled by measures taken after skeletal preparations and µCT scans of the various alleles (Fig. 2A). To verify that the ectopic gain of *Hoxb13* expression was indeed responsible for tail shortening, we analysed the two separate alleles (*HoxB^Del(i9-13):Hoxb13hd1^*) and *HoxB^Del(i9-13:Hoxb13hd2^*) where a secondary mutation inactivated HOXB13 *in cis* with the *HoxB^Del(i9-13)^* (Figs. 1A; 2A). Siblings from both breeding stocks were analysed for their tail length past 8 weeks of age and a clear rank of mean tail sizes emerged with *HoxB^Del(i9-13)^* homozygous tails shorter than *HoxB^Del(i9-13)^* heterozygous, shorter than wild type, shorter than *HoxB^Del(i9-13):Hoxb13hd1^* heterozygous, shorter than *HoxB^Del(i9-13):Hoxb13hd2^* heterozygous, shorter than *HoxB^Del(i9-13):Hoxb13hd1^* homozygous, shorter than *HoxB^Del(i9-13):Hoxb13hd2^* homozygous (Fig. 2B). As HOXB13 inactivation rescued the tail shortening effect, we concluded that reduced tail length directly resulted from the *Hoxb13* gain of function. As expected, there was no difference in tail length between the *HoxB^Del(i9-13):Hoxb13hd^*and the *Hoxb13^hd^* alleles (Fig. 2B, Fig. S2). All mice carrying a homozygous loss of function of *Hoxb13* had elongated tails, regardless of their complete genotype and in agreement with the initial study (9).

**Figure 2:**
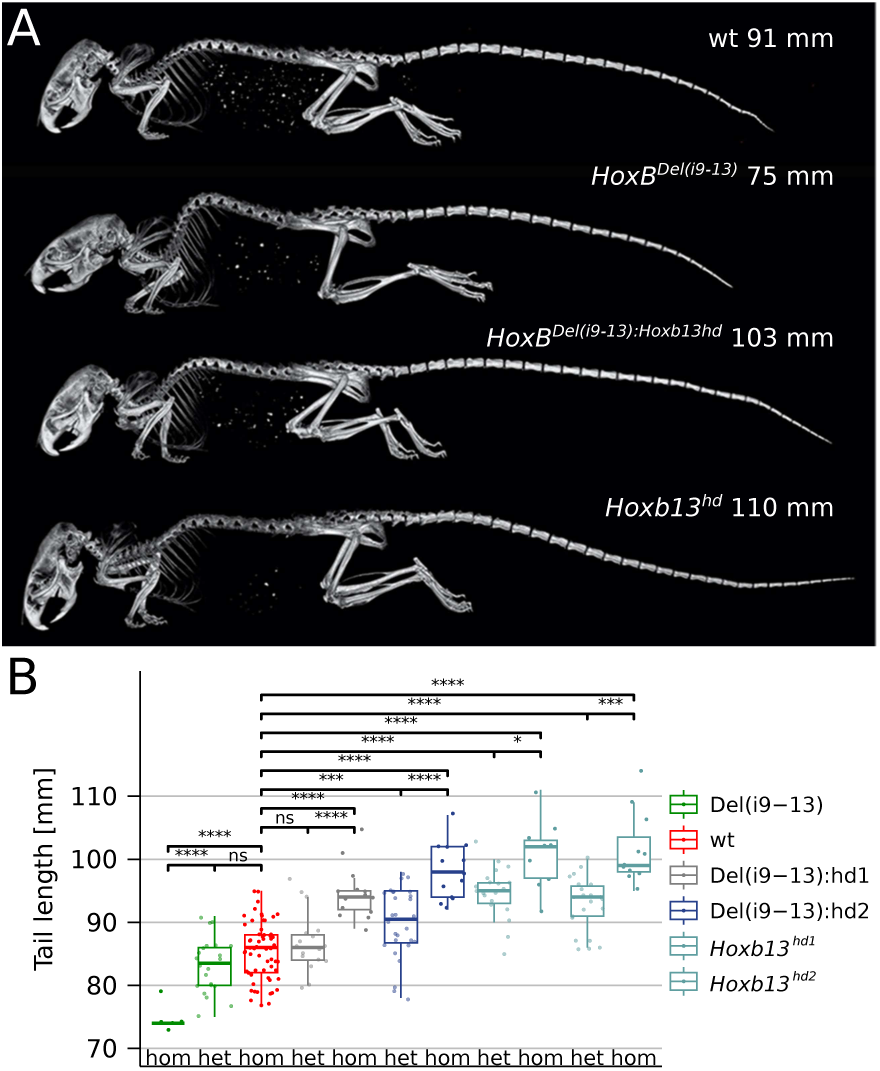
Effects of *Hoxb13* gain and loss of function upon tail length. (**A**) Lateral views of µCT 3D reconstructions of skeletons of representative phenotypes of adult animals carrying the various *HoxB* alleles shown in Fig. 1A. On top is a control animal with a tail length of 91 mm. Below is a *HoxB^Del(i9-13)^* homozygous specimen with a tail reduced to 75 mm (minus 18%). The inactivation of *Hoxd13 in-cis* with this deletion (below, *HoxB^Del(i9-13):Hoxb13hd^*) displayed a longer tail (103 mm, plus 13%), equivalent in length to those of mice lacking *Hoxb13* function (bottom). (**B**) Quantifications of tail lengths. While both homozygote *HoxB^Del(i9-13)^*(strong green) and *HoxB^Del(i9-13):Hoxb13hd^* (Del(i9-13):hd1, strong grey) and Del(i9-13):hd2 (strong blue) specimen displayed statistically significant deviations from controls (wt, red), their heterozygous versions (weak green and weak grey and blue, respectively) were much closer to wild type counterparts. hd1 and hd2 correspond to two independent lines. All wild-type littermates of all mutant lines were pooled together. Significance is assessed by the two-sided Welsh’s t-test (ns: p > 0.05, *: p <= 0.05, **: p <= 0.01, ***: p <= 0.001, ****: p <= 0.0001). Number of animals for each line (heterozygous and homozygous): Del(i9-13): 22 and 5, wt: 60, Del(i9-13):hd1: 17 and 14, Del(i9-13):hd2: 28 and 13, *Hoxb13^hd1^*: 20 and 9, *Hoxb13^hd2^*: 22 and 11. The lower and upper hinges correspond to the first and third quartiles. The upper whisker extends from the hinge to the largest value no further than 1.5 * IQR from the hinge.

We used skeletal preparations of adult specimen from the same breeding stock to evaluate to what extent such variations in tail length were due to changes in the number of vertebral types. Control specimen displayed between 28 and 30 complete caudal vertebrae (Fig. 3A). In Del(i9-13) homozygous, this number was between 24 and 27, while heterozygous displayed between 24 and 29 vertebrae (Fig. 3A). Skeletal alterations were also scored outside the tail, for all Del(i9-13) homozygous and most heterozygous displayed a number of ribs bearing thoracic vertebrae reduced by one (Fig. 3A, B), and a reduction in the number of lumbar vertebrae to L4 was sporadically observed (Fig. 3A, right). Therefore, the most affected individuals had a C7, T12, L4, S4, C25 vertebral formula, instead of the control C7, T13, L5, S4, C29 formula, prevalent in this background (Fig. S3). The defects in homozygous were more prevalent than in heterozygous and with higher expressivity, consistent with the gene dosage effect already observed in the tail length phenotype.

**Figure 3:**
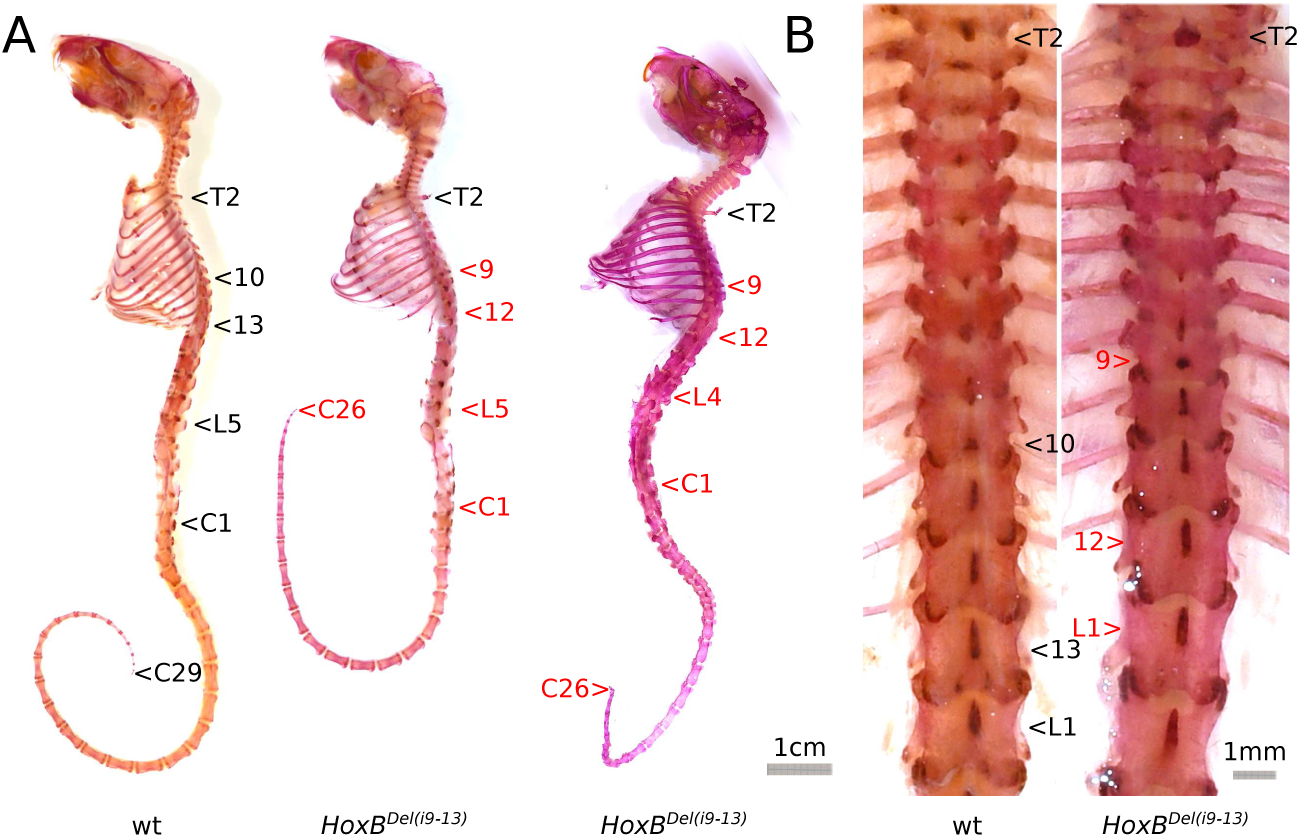
*Hoxb13* gain of function-dependent vertebral anomalies. (**A**) Alizarin-stained skeletal preparations showing the reproducible vertebral formulae in both wild type (left) and *HoxB^Del(i9-13)^* mutants (middle and right). The wild type skeletal pattern was composed of 7 cervical (C7), 13 thoracic (T13), 5 -rarely 6-lumbar (L5), 4 sacral (S4) and 29 -rarely 30-caudal (C29) vertebrae. The patterns most frequently observed in *HoxB^Del(i9-13)^* homozygous mutants are shown in the middle and right. **(B)** Magnification of the thoracic regions of a wt (left) and *HoxB^Del(i9-13)^* mutant (right). In several heterozygous and all homozygous mutants, T9 carried a neural spine typical of wt T10 (the so-called anticlinal vertebra). In such specimens, only 12 thoracic rib bearing vertebrae were scored and hence the T9-T10 exchange may be considered as a loss of normal T9 followed by a serial transformation. In the majority of hets and all homs, the caudal series was composed of less than 29 complete vertebrae. In the depicted mutant specimens, the number was 26, instead of 29. The corresponding quantifications are in Fig. S2B.

This dosage effect was further assessed by adding to Del(i9-13) animals a *Hoxd13* gain of function produced by the deletion of *Hoxd10* to *Hoxd12* included, i.e., a *HoxD* allele identical to the Del(i9-13) allele on the *HoxB* cluster, bringing *Hoxd13* next to *Hoxd9*. In this case, a transient gain of *Hoxd13* expression was observed (25). *HoxB^Del(i9-1)^*:*HoxD^Del^*(*^10–12^*) compound mutants showed a further decrease in the number of caudal vertebra when compared to *HoxB^Del(i9-13)^* (Fig. S2B). This decrease was minor yet statistically significant. We concluded that group 13 genes might share a function in limiting posterior growth zone elongation during generation of tail somites. However, the contribution of *Hoxb13* in this task was clearly more prominent than that of *Hoxd13*.

### Transcriptome analyses

We looked at the impact of *Hoxb13* gain of function on the embryonic transcriptomes at E9 and E10. RNA was extracted from ‘posterior embryos’, i.e., dissected from below the forelimb buds, and three samples of each genotype were used. *Hoxb13* transcript levels were dramatically increased up to 17 FPKM in E9 Del(i9-13) samples (adjusted p-value of 1e-66), as well as to ca. 4 FPKM in both E9.5 and E10.5 *HoxB^Del(i9-13)^* and *HoxB^Del(i9-13):Hoxb13hd^* (adjusted p-value of 0.02 and 0.004 respectively; Fig. S4A). Principal component analysis confirmed that most of the variance was likely due to the developmental stage (Fig. S4B). The second principal component spread the samples according to genotypes. The relative position of the *HoxB^Del(i9-13)^* transcriptome was concordant between both stages and the *HoxB^Del(i9-13):Hoxb13hd^* samples were expectedly more similar to control. Expression levels of all *Hox* genes confirmed that the assigned developmental stages were indeed correct and that no major change was scored beside *Hoxb13* RNAs (Fig. S4C).

We looked for RNAs which were modulated between wild-type and *HoxB^Del(i9-13)^* embryos at both stages, and inversely regulated in the *HoxB^Del(i9-13):Hoxb13hd^* allele. We identified only two genes significantly up-regulated with *Hoxb13* gain of function (*Chl1*, *Baiap2l1,* Fig. S5A), whereas a single gene was down-regulated (*Rbpj*, Fig. S5B). The decrease in *Rbpj* mRNAs is noteworthy, for *Rbpj* loss of function mutants show a developmental arrest during early somitogenesis (27).

### ES cells-derived gastruloids as proxies to study *Hoxb13* insulation

To study the importance of the DNA spacer length *versus* the presence and number of CBSs, in the necessary insulation of *Hoxb13,* we turned to gastruloids, i.e., mES cells-derived pseudo-embryos (28, 29), which are excellent proxies to study the extending posterior part of mammalian embryos (30, 31). Gastruloids can be produced in large amount and are thus amenable to high throughput analyses (18). We reproduced in ES cells the same Del(i9-13) deficiency and produced heterozygous gastruloids. We examined the expression of both *Hoxb9* and *Hoxb13* by WISH in control and mutant gastruloids and found no salient difference in the expression of *Hoxb9* at 120 hours after aggregation (AA) (Fig. 4A). In both cases, the entire extending (posterior) part was positive with a clearly delimited spatial boundary at a more ‘anterior’ level. At this stage, as well as at 144h AA, control gastruloids did not show any trace of *Hoxb13* mRNAs, yet a strong signal was detected in Del(i9-13) mutant specimen, with a full penetrance (Fig. 4A, B). The position of the expression boundary was slightly more posterior than that of *Hoxd9* (Fig. 4C) in agreement with the colinear distribution of *Hox* transcripts observed in gastruloids (18, 30).

**Figure 4:**
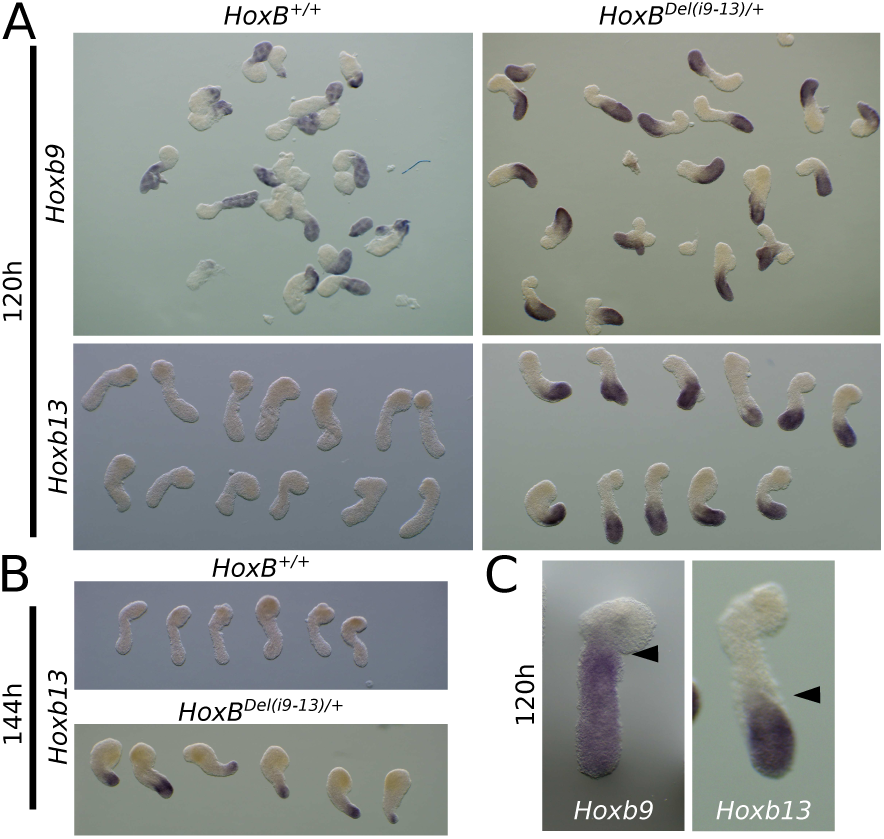
*Hoxb9* and *Hoxb13* expression in control and mutant gastruloids. (**A**) Whole-mount in situ hybridisation for *Hoxb9* top and *Hoxb13* bottom on either wild-type (left) or *HoxB^Del(i9-13)/+^* (right) 120h gastruloids showing the clear and penetrant gain of *Hoxb13* in the mutant condition. (**B**) WISH of *Hoxb13* on wild-type (top) and *HoxB^Del(i9-13)/+^* (bottom) 144h gastruloids. (**C**) Comparison of the pattern of expression of *Hoxb9* in wild-type and *Hoxb13* in HoxB^Del(i9-13)/+^ gastruloids at 120h, showing colinearity in the anterior-posterior (AP) expression border (arrowheads).

### Same cellular population in both control and *Hoxb13* gastruloids

To look at which cells precisely expressed the gained *Hoxb13* mRNAs, as well as to evaluate any potential effects of this gain of function either upon gene expression, or on the distribution of cell types in these mutant gastruloids, we carried our single cell RNA analysis using 144h gastruloids, a stage where *Hoxb13* expression was well established in mutant specimens. The analysis of this scRNAseq dataset revealed that the distribution of cellular clusters remained virtually unchanged between control and mutant gastruloids (Fig. S6A). It also showed that gained *Hoxb13* mRNAs were mostly found within neuro-mesodermal progenitors (NMP) cells, as well as within the ‘Neural Tube 1’ cluster, likely composed of early differentiating neuronal cells, adjacent to the NMP cluster (Fig. S6B). Gene clustering based on all *Hox* gene expression throughout cell fates revealed that *Hoxb13*, which normally clusters with its paralogous *Hoxa13* gene, would now cluster with group 9-10 *Hox* genes in the mutant gastruloids, i.e., *Hoxb13* would be expressed in cells with a general context related to the latter genes (Fig. S6C, arrowheads).

Using baredSC (32), the exact co-expression of genes was assessed and, while no positive correlation was found with the expression of either *Hoxa9*, *Hoxc9* and *Hoxd9*, a strong positive correlation was observed with the expression of *Hoxb9*, i.e., the new immediate neighbour of *Hoxb13* after deletion of the spacer DNA. Indeed, 54 percent (+/- 14%) of cells positive for *Hoxb9* were also positive for the ectopic *Hoxb13* mRNAs (Fig. S6D, E), illustrating that the relocated *Hoxb13* gene was transcribed at the time and in the cells where an elusive *Hoxb10* gene would be transcribed, should it still be present in the amniote *HoxB* cluster, as is the case for some anamniotes species like zebrafishes. This illustrates once more the decisive role played by the relative position of *Hox* genes in their respective cluster, rather than by their promoters, in the precision of their transcriptional regulation (refs in (33).

Altogether, the gain of *Hoxb13* expression did not drastically modify the transcription landscape of gastruloids at 120h, when the analysis was carried out using control and mutant specimens from the same batch. This observation is coherent with the comparable analysis in mouse embryos (see above) and further confirmed that the effect of this gain of function is mostly quantitative, i.e., involving differences in transcription timing, rather than purely qualitative such as starting a distinct transcription program, in agreement with the homeotic phenotypes observed in mice.

### Insulation of *Hoxb13* from the *HoxB* cluster

Next, we carried out a series of analyses in 132h gastruloids to better document the role of this spacer DNA in the isolation of *Hoxb13* from earlier and more ‘anterior’ regulations. At this stage, the H3K27 acetylation (H3K27ac) not only covered the entire *Hoxb1* to *Hoxb9* region, but also included the nearby located *Mir196a-1* locus and the *GM53* LncRNA, i.e., a region including both CBS5 and CBS6 (Fig. 5A), indicating that this immediate neighbourhood was also actively transcribed. The ChIP profile of NIPBL, a factor that helps loading the cohesin complex, revealed an enrichment throughout this H3K27ac positive region, which matched with the distribution of RAD21, a sub-unit of the cohesin complex that was enriched all over this region, with a strong accumulation at CBS7 and CBS8 (Fig. 5A). From this we concluded that, as in the case of the *HoxD* cluster (18), cohesin deposition and loop extrusion occurred in an asymmetric manner. In this case, cohesin was already detected passed the *Hoxb9* position and hence the main blockage to looping up to *Hoxb13* was likely achieved by either CBS7 or CBS8, or both, despite their opposite orientations.

**Figure 5:**
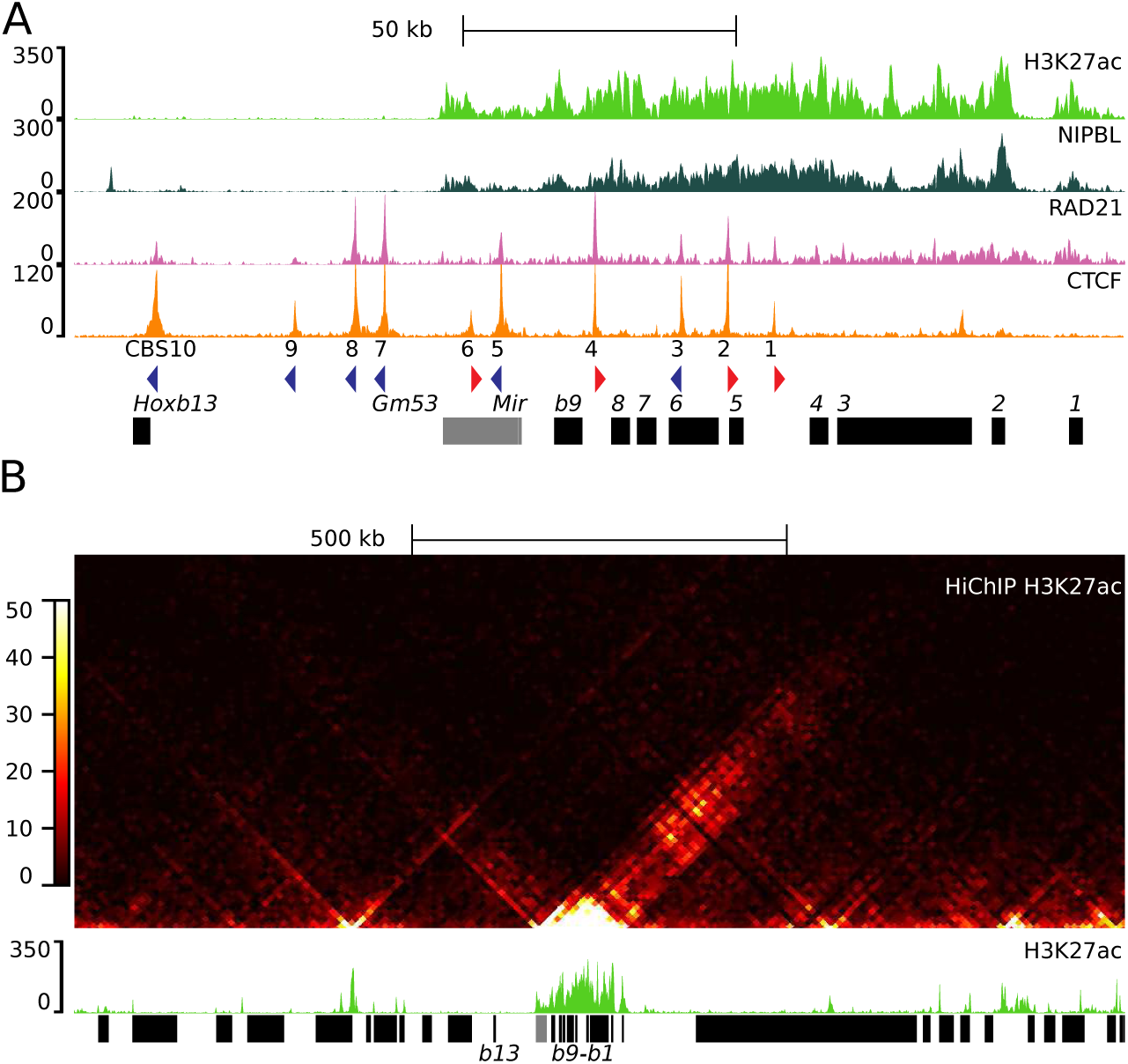
Chromatin structure of the *HoxB* locus in late gastruloids. (**A**) Profiles of either H3K27ac (light green), NIPBL (dark green) and RAD21 (magenta) in 132h gastruloids, or CTCF at 168h over the *HoxB* locus (data extracted from (18), GSE205779). CTCF binding sites (CBS) are numbered 1 to 10 from *Hoxb1* to *Hoxb13* and the coloured arrowheads below indicate orientations. *Hoxb* genes are shown as black rectangles below. The *HoxB* cluster is active until the *Mir196a-1* gene, while *Hoxb13* is tightly isolated (**B**) HiChIP of H3K27ac over a Mb of DNA containing the *HoxB* cluster (indicated below), with the matching profile of H3K27ac ChIP-seq of 120h gastruloids below (green) (data extracted from (18), GSM6226299 and GSM6226246). Black rectangles below represent protein coding genes. *Gm53* and *Mir196a-1* are in grey. The vast majority of contact between ‘active’ regions, i.e., involving H3K27ac, are towards the TAD located 3’ of the cluster, whereas *Hoxb13* is not at all involved.

To verify this directionality in regulation, we analysed a H3K27ac HiChIP dataset, which revealed that this entire transcribed region strongly interacted with sequences located exclusively in 3’, i.e., within the same TAD, confirming that the early activation of *Hox* genes mostly involves regulatory inputs coming from this TAD (18, 34), whereas the ‘posterior’ TAD, including *Hoxb13*, was tightly insulated from these regulations with no signal involving *Hoxb13* itself (Fig. 5B).

To further assess the mechanism of *Hoxb13* insulation, in particular to discriminate between the effect of the linear genomic distance *versus* the number and orientation of CTCF sites, we produced an ES cell clone deficient for one copy of the entire *HoxB* locus, i.e., carrying a targeted deletion including from *Hoxb1* to *Hoxb13* (*HoxB^Del^*; Fig. 6A). A deletion of the DNA spacer was then induced on the other chromosome, thus leading to the *HoxB^Del/Del(i9-13)^* configuration (Fig. 6). Similar to the *HoxB^Del(i9-13)/+^*heterozygous gastruloids, their *HoxB^Del/Del(i9-13)^* mutant counterparts expressed *Hoxb13* prematurely and at an ectopic anterior position in 120h gastruloids (Fig. 6C), whereas *Hoxb9* expression remained unchanged (Fig. S7). Noteworthy, *HoxB^Del/Del(i9-13)^* mutant gastruloids displayed some variability (up to three folds) in the strength of *Hoxb13* activation between different replicates (Fig. S7).

**Figure 6:**
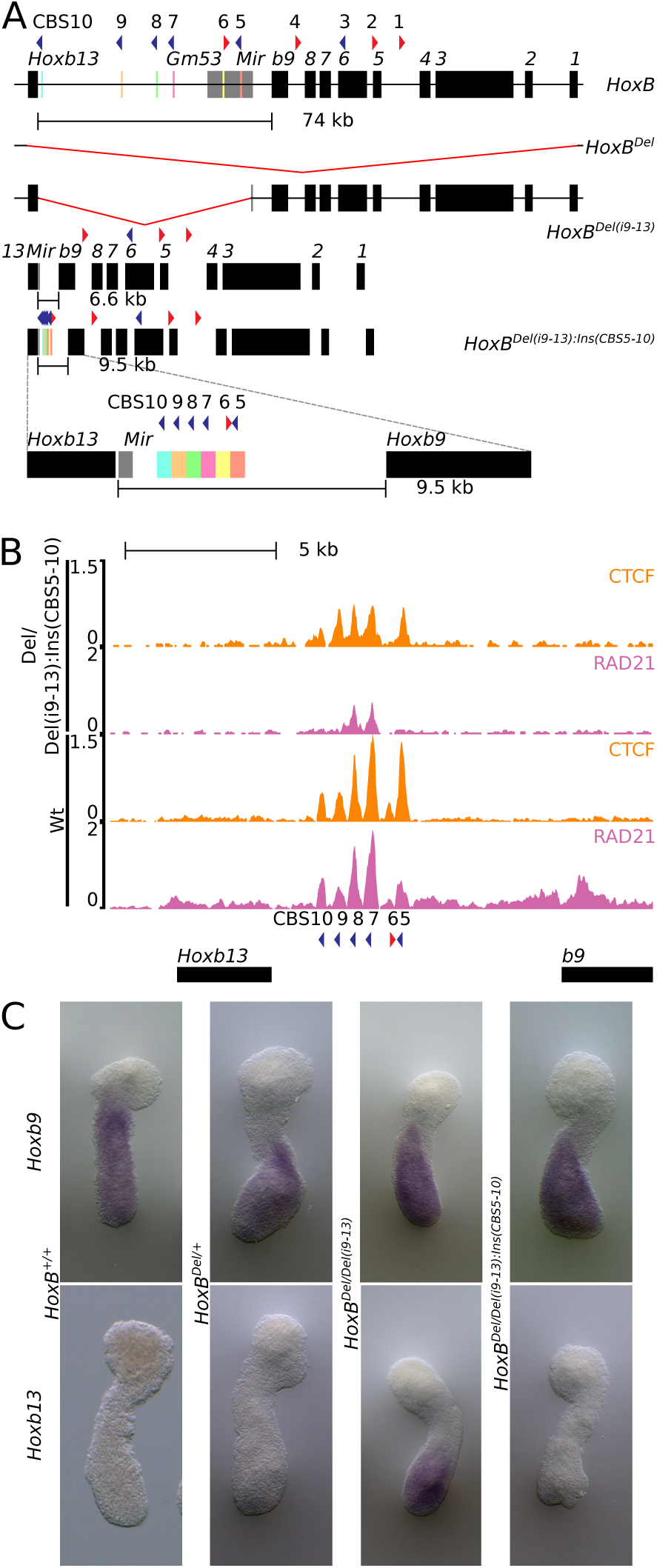
Insertion of a CTCF cassette into the deleted spacer DNA. (**A**) Schematic of the various *HoxB* alleles with on top the wild type *HoxB* locus along with the 74 kb large DNA spacer indicated as a black line. Below is shown the large deletion of the entire *HoxB* locus used to balance the other alleles. The third track shows the deletion of the (9–13) intergenic region, leading to the ‘consolidated’ *HoxB* cluster shown in the fourth track. In this latter configuration, the distance between *Hoxb9* and *Hoxb13* is now 6.6 kb without any CTCF binding sites. The fifth track shows the allele after insertion of a cassette containing the 6 native CBS separated from one another by ca. 0.5 kb. The distance between *Hoxb9* and *Hoxb13* is now of 9.5 kb. This new intergenic region is magnified below, with each coloured block containing a CBS corresponding to those of the wild-type allele, in the right order (see first track). (**B**) ChIP-seq profiles of CTCF (orange) and RAD21 (magenta) using the *HoxB^Del/Del(i9-13):Ins(CBS5-10)^* allele (the insertion of the cassette) on top and wt on bottom on 96h gastruloids aligned on the *in silico* reconstructed Del(i9-13):Ins(CBS5-10) mutant genome. (**C**) Whole mount *in situ* hybridisation with *Hoxb9* (top) and *Hoxb13* (bottom) probes on 120h gastruloids with the following genotypes from left to right: *HoxB^+/+^, HoxB^Del/+^, HoxB^Del/Del(i9-13)^* and *HoxB^Del/Del(i9-13):Ins(CBS5-10)^.* The insertion of the CTCF cassette fully suppresses the *Hoxb13* gain of function, even with a much shorter distance between *Hoxb9* and *Hoxb13*. For quantifications, see Fig. S7.

We used the *HoxB^Del/Del(i9-13)^* ES cells as hosts to reintroduce the complete series of deleted CBSs, yet without the large DNA spacing in between. We used a synthetic DNA cassette containing the six CBSs initially identified in the spacer DNA (CBS 5 to CBS 10, Figs. 1, 6), but separated from one another by ca. 500 bp. The overall size of this cassette was thus of ca. 3 kb, instead of the 70 kb length that normally separates *Hoxb9* from *Hoxb13*. Homology arms were introduced and the cassette was recombined between *Hoxb9* and *Hoxb13*, on top of the *HoxB^Del(i9-13)^* chromosome to produce the *HoxB^Del/Del(i9-13):Ins(CBS5-10)^* allele. The cassette contained all six CBSs in their native orientations and hence the number, orientation and relative sequence of CBSs were as in the wild type chromosome, except that the genomic distance was reduced to a size generally found between any neighbouring *Hox* genes (Fig. 6A, bottom).

ChIPmentation (ChIPM-seq) was carried out using 96h mutant gastruloids to check for the presence of both bound CTCF and the cohesin complex through its RAD21 subunit. Sequence reads were aligned onto an *in silico* reconstructed Del(i9-13):Ins(CBS5-10) mutant genome. The CTCF profile observed was comparable to the control situation in terms of relative intensities, with CBS5, 7 and 8 being prominent whereas CBS6 was barely scored (Fig. 6B, compare first and third profiles). Likewise, RAD21 mostly accumulated at CBS7 and CBS8, as in the control situation (Fig. 6B; second and fourth profiles), even though the global accumulation was weaker, in part due to the hemizygosity of the locus. The recombination of this CBS cassette between *Hoxb9* and *Hoxb13* in ES cells had no detectable effect upon the mRNA levels of either *Hoxb1* or *Hoxb9* in gastruloids derived thereof (Fig. 6C, top). In contrast, gastruloids produced out of *HoxB^Del/Del(i9-13):Ins(CBS5-10)^* ES cells had no detectable expression of *Hoxb13* by WISH and mRNA levels were poorly significant, either in 120h gastruloids, or later at 144h (Fig. 6C, bottom; Fig. S7).

To further verify that the insertion of the cassette in this *HoxB^Del/Del(i9-13):Ins(CBS5-10)^* allele had not structurally affected the *Hoxb13* transcription unit in any way, the mutant locus was entirely DNA-sequenced by nCATS (35) and no obvious off-target modifications were scored (Fig. S8A, B). Furthermore, we deleted back the cassette after its insertion (the *HoxB^Del/Del(i9-13):Ins(CBS5-10:Del(CBS5-10)^* _and RNA-seq experiments comparing the *HoxB*_*^Del(i9-13)^*_mutant gastruloids at 120h with the *HoxB*_*^Del/Del(i9-13):Ins(CBS5-10)^* _and the *HoxB*_*^Del/Del(i9-13):Ins(CBS5 10:Del(CBS5-10)^* versions (with and without the CTCF cassette inserted) were carried out. Deletion of the cassette led to an upregulation of *Hoxb13* transcription close -if not equal (65%)- to that observed before its recombination (Fig. S8C). From this, we concluded that this synthetic cassette was capable to achieve most of *Hoxb13* insulation in gastruloids, despite the short distance between *Hoxb9* and *Hoxb13* in the *HoxB^Del/Del(i9-13):Ins(CBS5-10)^* mutant allele.

### CTCF and *Hoxb13* insulation

While the CTCF ChIPM-seq profiles of both the control and the *HoxB^Del/Del(i9-13):Ins(CBS5-10)^*mutant allele indicated that all CBS can be bound (Fig. 6B), it did not reveal whether all (or several) sites must be bound simultaneously to achieve full insulation or, alternatively, if CTCF binding to only few sites is sufficient. We thus deleted either three or four consecutive CBSs from the cassette (Fig. 7) and look at the effect upon *Hoxb13* transcription both by WISH and RNA-seq. ES cells carrying these deletions were verified for the presence of bound CTCF on the remaining sites either by ChIPmentation or by CUT&RUN approaches (Fig. 7A). While the deletion of CBS8 to 10 (Fig. 7A; Del(CBS8-10)) did not elicit any detectable up-regulation of *Hoxb13* (Fig. 7B), a slight though significant increase was scored when the four CBS7 to 10 were deleted (Fig. 7A, B; Del(CBS7-10)). The latter increase was nevertheless detectable by WISH as a weak but clear signal in most posterior gastruloid cells (Fig. 7C, arrowhead). However, as was the case for the *HoxB^Del/Del(i9-13)^*gastruloids, the response to this partial CBS deletion was somehow variable amongst replicates and data in Fig. 7C show gastruloids with the highest amount of *Hoxb13* mRNAs. These results indicated that, while only 2 CBSs with opposite orientations could not achieve full isolation, they were still capable to importantly delay ectopic *Hoxb13* activation.

**Figure 7:**
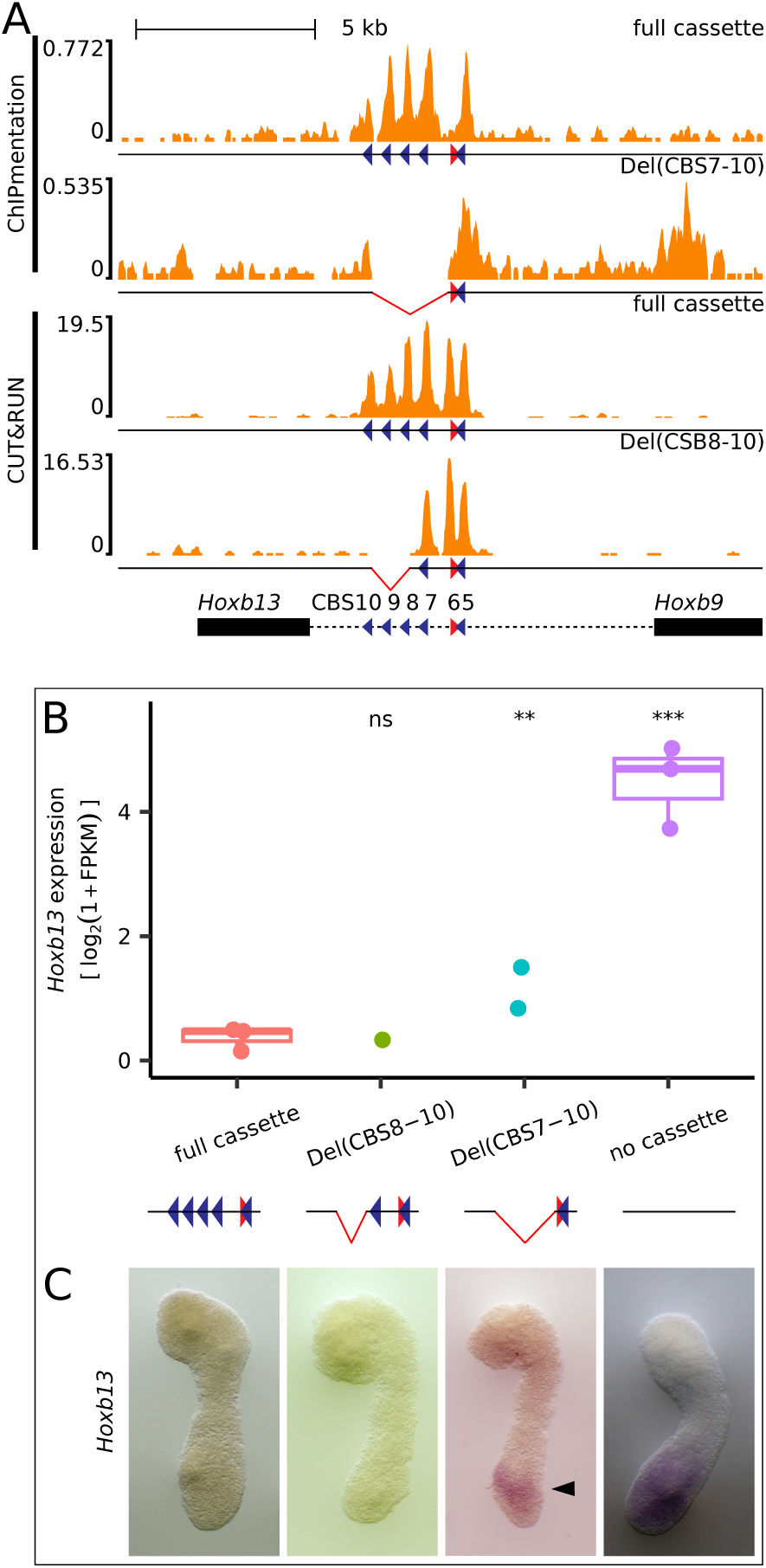
Deletion of CTCF binding sites from the inserted CTCF cassette. (**A**) ChIPmentation on 96h gastruloids (top two profiles) and CUT&RUN on ES cells (two profiles on bottom) of CTCF from the *HoxB^Del/Del(i9-13):Ins(CBS5-10):Del(CBS7-10)^*(top, Del(CBS7-10)) and *HoxB^Del/Del(i9-13):Ins(CBS5-10):Del(CBS8-10)^*(bottom, Del(CBS8-10). (**B**) Quantification of *Hoxb13* mRNAs in the two deletions shown in **A**. While in Del(CBS8-10) *Hoxb13* is still tightly isolated, some transcripts start to appear in the Del(CBS7-10). The significance was assessed by the Wald test on counts with DESeq2 in comparison to the full cassette condition. The p-values were not corrected for multiple tests: ns: p > 0.05, *: p <= 0.05, **: p <= 0.01, ***: p <= 0.001. (**C**) WISH of *Hoxb13* transcripts in the two deletions shown under A. A weak signal appears at the right position in the Del(CBS7-10) gastruloids (black arrowhead), not detected in the Del(CBS8-10) counterparts.

Finally, we looked at the insulating capacity of the native CBSs re-introduced into the *HoxB^Del/Del(i9-13)^* chromosome though with inverted orientations. The inversion of the full re-inserted cassette (*HoxB^Del/Del(i9-13):Ins(CBS5-10):Inv(CBS5-10)^*) had an insulating capacity close to -if not identical with- the insertion of the cassette carrying the CBSs in their native orientation, suggesting that the orientation of the CTCF sites was not the main determinant of the insulation, at least in this particular topology (Fig. S9). We also produced a *HoxB^Del/Del(i9-13):Ins(CBS5-10):Del(CBS7-10)^* allele where the remaining CBS5 and CBS6 had an inverted configuration. Indeed, these two sites displayed different binding behaviours, with CBS5 strongly bound by CTCF and accumulating RAD21, whereas its close neighbour CBS6 poorly binding CTCF. While gastruloids produced from this *HoxB^Del/Del(i9-13):Ins(CBS5-10):Inv(CBS5-10):Del(CBS7-10)^*) configuration showed a gain of *Hoxb13* expression (Fig. S9A, B), the leakage in insulation was in the same range than that observed before the inversion of these two CBSs.

## DISCUSSION

Past experimental approaches have suggested that *Hox* genes belonging to the paralogy group 13 control the end of axial extension during vertebrate development. Accordingly, their functional modulation could be determinant in species-specific body lengths (9–11). In addition, adaptive variations in tail lengths under natural conditions were associated with few QTLs in deer mice, amongst which one affecting *Hoxd13*. In this case, a lower amount of *Hoxd13* was associated with a larger presomitic mesoderm (PSM), from where somites derive, due to increased number of NMP cells (4). Furthermore, *Hoxb13* was recently identified as the main candidate gene for tail length in Chinese long-tailed sheep breeds (14). Here we demonstrate that a physiological gain of function of *Hoxb13* leads to both a drastic variation in tail length and other homeotic transformations at more rostral body levels. Such phenotypes associated with the ectopic expression of *Hox13* genes could explain the evolution of a TAD boundary always located between the latter genes and the rest of the gene clusters thereby insulating this terminal function and delaying its implementation until the appropriate time has come (20). Despite its particular structure lacking most other posterior *Hox* genes, a strong and constitutive TAD boundary was maintained in *HoxB* (23) coinciding with both a large DNA spacer and the preservation of a number of CTCF sites, as in the three other *Hox* gene clusters (18).

### A physiological gain of function causing a heterochronic phenotype

Removal of the DNA region between *Hoxb9* and *Hoxb13* induced premature anterior expression of *Hoxb13*, leading to vertebral column shortening, including vertebra loss from the lower thoracic, lumbar and caudal regions. These defects were all rescued when the ectopic gained HOXB13 protein was inactivated *in cis*, demonstrating causality. These phenotypes were much weaker than those produced by using ‘transgenic’ gain of function where neither the amount of ectopic protein, nor its precise timing of delivery can be precisely controlled (10, 36). In contrast, the extent of the morphological anomalies reported here is both dose dependent and in the range of what was previously reported using *on-site* genomic modifications for *Hox13* genes (e.g., (9, 37)). Therefore, the dramatic developmental patterning defects observed upon ectopic expression of *Hox13* genes (6, 7, 36), while illustrating a genuine effect should be considered as an extreme outcome of overloading the system with HOX13 proteins, rather than as a genuine physiological response to a slight variation in *Hox13* transcripts as it may naturally occur.

The deletion of the DNA spacer re-positioned *Hoxb13* near *Hoxb9*, at the relative place of an elusive *Hoxb10*, absent from the amniote *HoxB* cluster, though still present in teleost fishes. As a consequence (38, 39), *Hoxb13* transcription was activated prematurely and its gain of expression was scored as early as E9 in the posterior growth zone. By late E10, signal intensity dropped significantly, resulting in a short window of *Hoxb13* exposure during somitogenesis. This was nevertheless enough to impact the morphology of the vertebral column at a level far more rostral than expected for *Hoxb13*, up to the ninth thoracic vertebra, reflecting an ectopic transitory expression in progenitors of somite 20 and more posterior. This thoracic 9^th^ rostral limit of the *HoxB^Del(i9-13)^* phenotypes is of interest since in the absence of the *Hoxb1-Hoxb9* part of the cluster (*HoxB^Del^*(*^1–9^*)), which left the *HoxB(i9-13)* spacer region untouched and thus likely preserved the insulation of *Hoxb13* from the influence of anterior enhancers, malformations were limited to thoracic vertebra 8 or more anterior (40). We conclude that the presence of the spacer region protects the lower thoracic, lumbar and sacral midsegment of the main body axis from a growth arrest by preventing premature *Hoxb13* expression in the posterior growth zone.

The phenotypes observed in the lumbar and the tail regions are notable, for L4 or L3 types of lumbar vertebral formulae had previously been reported only in very few stocks of laboratory mice, either in animals carrying the *Dominant hemimelia* (*Dh*) mutation (41), or in mice with premature expression of posterior *Hox* transgenes (42). Regarding tail length, the gain of function, rescue and loss of function effects indicate that *Hoxb13* alone could account for at least one third of length variation of the adult tail in laboratory mice. Premature presence of HOXB13 made tails shorter and the absence of *Hoxb13* homeobox made tails longer, both in a gene dose dependent manner. Combined gain of function mutations between *Hoxb13* and *Hoxd13* showed only a subtle increase in tail shortening despite recent results in deer mice where *Hoxd13* seems to be prominent (4).

Tail bud NMP cells display progenitor activity partly relying on *Hox13*, *Gdf11* and *lin28* genes (6) and forced expression of *Lin28b* increased tail vertebra count by as much as five (7), a hypermorphosis that corresponds to the effect of *Hoxb13* inactivation. Conversely, both the hereby described *Hoxb13* gain of function and the *Lin28a* and *Lin28b* double knockouts reduced vertebra count by approximately four, suggesting a window of ca. 10 vertebrae where this system can operate. We did not detect any significant change in *Lin28a* or *Lin28b* transcript levels at E10 in our *Hoxb13* loss- or gain-of-function, which is consistent with *Hoxb13* being downstream a *Lin28/let-7* driven genetic regulatory circuit.

### Transcriptomes analyses and Insulation by CTCF sites

The gain of *Hoxb13* function had a moderate effect on global gene transcription, with only a handful of genes up- or down regulated. This was not unexpected when dealing with a heterochronic shift in an otherwise normal process, i.e., the termination of tail extension, *a fortiori* under physiological conditions. However, the down-regulation of *Rbpj* is of interest, for transcripts accumulation in the tail of this effector of the *Notch* pathway is reminiscent of *Hoxb13* (Fig. S3A, B, C). Homozygous *Rbpj* loss of function mutants have severe arrest of early somitogenesis (27), which makes it possible that *Rbpj* mediates the effect of HOXB13 during vertebral column development, in both normal and ectopic contexts. Such potential interactions between *Hox* genes and the *Notch* pathway had been previously proposed (43).

Using gastruloids, we show that cells expressing ectopic *Hoxb13* mRNAs are a subset of those expressing *Hoxb*9 within the NMP population, further supporting the claim that *Hox* genes are expressed in time and space along the AP axis mostly in response to their relative position within the clusters rather than based on specific regulatory sequences (44). We also show that a small CTCF cassette re-established most of the *Hoxb13* insulation, at least until the end of our gastruloid cultures (168h) and hence that these sites are necessary and even sufficient in these conditions for tight insulation to occur. When the cassette was reduced to three or two CTCF sites, a leakage occurred suggesting that the reiteration of CBS in this spacer region was important. Recent work has shown that during gastruloid development, loop extrusion initially occurs asymmetrically upstream the *Hox* clusters (on the side of *Hox1*), as suggested by an enriched deposition of both RAD21 and NIPBL (18). In this view, these CBSs may represent as many potential blocks to loop extrusion, inducing transcription delays in an additive manner before reaching *Hoxb13*.

Two transcription units were nevertheless found right in 5’ of *Hoxb9* (*GM53* and *Mir196a-1*), which were also enriched in both cohesin and NIPBL thus adding 19 kb of chromatin potentially active in loop extrusion upstream *Hoxb9* and including both CBS5 and CBS6. Therefore, loop extrusion initiated from this part of the ‘spacer’ DNA would be blocked or delayed mostly by CBS7 and CBS8. The orientation of these CBSs did not seem to be critical in their insulation capacity, either when the full cassette was inverted, or when only two occupied sites were left in either orientation, in agreement with previous work ((19) and ref. therein).

If the insulation of *Hoxb13* mostly depend upon the presence of CBSs, then one may wonder why this large DNA spacer still exists in all vertebrates *HoxB* clusters examined thus far. The probability of reducing the length by spontaneous internal deletions not involving any CTCF site may be low, despite an enrichment in repeats (SINE, LINE, LTR) when compared to the rest of the cluster, which as all *Hox* clusters are generally depleted of such sequences (45). Another possibility is that other DNA sequences present in this spacer may play a role during development or adulthood related or unrelated to the ‘classical’ function of *Hox* genes, which evolutionarily constrained this DNA interval. The presence of some DNA sequences conserved within all mammals may support this hypothesis, even though two of them precisely correspond to CBS7 and CBS8.

## ACKNOWLEDGEMENTS

We thank all the colleagues from the Duboule laboratories for discussions, Dr. Christopher C. Bolt for his help and Anne-Catherine Cossy for technical assistance. This work was supported in part using the resources and services for sequencing of the Gene Expression Research Core Facility (GECF) at the School of Life Sciences of EPFL, and the Geneva iGE3 Genomics Platform (University of Geneva). In addition, all computational work was performed using the SCITAS facilities of the Scientific IT and Application Support Center of EPFL. This work was supported by funds from the École Polytechnique Fédérale de Lausanne (EPFL, Lausanne), the European Research Council grant RegulHox (588029) and the Swiss National Science Foundation (CRSII5_189956 and 310030B_138662).

## MATERIALS AND METHODS

### Animals

All experiments involving mice were approved and performed in compliance with the Swiss Law on Animal Protection (LPA) under license numbers GE 81/14 and VD2306.2 (to D.D.). All animals were kept in a continuous back cross with C57BL6 × CBA F1 hybrids. Mice were housed at the University of Geneva Sciences III animal colony with light cycles between 07:00 and 19:00 in the summer and 06:00 and 18:00 in winter, with ambient temperatures maintained between 22 and 23 °C and 45 and 55% humidity, the air was renewed 17 times per hour.

### Generation of alleles

We generated the *HoxB^Del(i9-13)^* allele by pronuclear injection of a mixture of two plasmids each containing the coding region for the Cas9 enzyme and a selected template for single guide RNA (Table S1). The deficiency included 67403 bp (chr11:96,197,567-96,264,470 mm10) in the *HoxB* cluster. Zygotes fertilized by Del(i9-13) homozygous mutant males and wild-type females were injected with a mix of two plasmids expressing specific guide RNAs corresponding to two nearby sequences flanking two thirds of the homeodomain of *Hoxb13* (chr11:96,196,015 and chr11:96,196,172) (Table S1). Animals were genotyped by PCR (Table S2). In this way, we produced four different alleles where the *Hoxb13hd* mutant was isolated either *in cis* with *HoxB^Del(i9-13)^* (the *HoxB^Del(i9-13):Hoxb13hd1^* and *HoxB^Del(i9-13):Hoxb13hd2^* alleles), or *in trans*, i.e., on the wild type chromosome (the *Hoxb13^hd1^*and *Hoxb13^hd2^* alleles). The sequences of both the various breakpoints and the mutated *Hoxb13* homeodomains are in Tables S3 and S4, respectively. All four founders were crossed with wild type breeders in three consecutive generations to segregate away potential CRISPRcas9 off-target events.

### Other mutant mice

The mutant strains *HoxD^Del^*(*^10–12^*) and *Hoxd13^hd^* were previously published (25, 26).

### WISH

Whole mount *in situ* of mouse embryos were performed as described in (46), using probes in Table S5. Image acquisition was as in (43).

### Skeletal preparations

Skeletal preparations were carried according to conventional methods (47). After Alizarin redS staining, the specimens were dehydrated into concentrated glycerol and photographed with a Nikon D810 camera, equipped with AF-S Micro NIKKOR 60mm f/2.8G ED objective and Nikon SB-910 SPEEDLIGHT.

### Micron-scale computed tomography (µCT)

Measurements of the vertebrae were carried out in the OsiriX MD v.10.0.1 software, in the 2D orthogonal MPR views of 3D reconstructions using WL/WW CT-Bone mode, displayed at 0.1 mm thick slab setting. Each measurement was taken when the object was approximately zoomed to five centimeters on the screen in one of the three windows, and the measure was manually defined using the length tool over the bone section. In all three windows, the longest head to tail length of all vertebrae were scrolled to, recorded and the longest of the three values were used. Jpg images and movies were exported at preset modes.

### Tail length measurements

For the tail length growth trajectories (Fig. S2A), data were pooled from five litters of two separate stocks born from heterozygous parents. Eleven and nine homozygous individuals carrying either the *Hoxb13^hd1^*or the *Hoxb13^hd2^* alleles, respectively, and a total of fifteen wild type littermates were documented two to three times weekly, by taking direct measures of the length between the anus and the tip of the tail on a millimeter scale. The same measurement method was used than in Fig. 2B, but in Fig. 2B, each dot represents a different animal while in Fig. S2B each dot is a measurement.

### Production of gastruloids

Mouse embryonic stem (mES) cells were routinely cultured in gelatinized tissue-culture dishes with 2i (48). LIF DMEM medium composed of DMEM + GlutaMAX supplemented with 10% ES certified FBS, non-essential amino acids, sodium pyruvate, beta-mercaptoethanol, penicillin/streptomycin, 100 ng/ml of mouse LIF, 3 µM of GSK-3 inhibitor (CHIR99021) and 1 µM of MEK1/2 inhibitor (PD0325901). Cells were passaged every 3 days and maintained in a humidified incubator (5% CO2, 37 °C). The differentiation protocol for gastruloids was previously described (30). ES cells were collected after accutase treatment, washed and resuspended in prewarmed N2B27 medium (50% DMEM/F12 and 50% Neurobasal supplemented with 0.5x N2 and 0.5x B27, and with non-essential amino acids, sodium pyruvate, beta-mercaptoethanol, penicillin/streptomycin). 300 cells were seeded in 40 µl of N2B27 medium in each well of a low-attachment, rounded-bottom 96-well plate. 48h after aggregation, 150 µl of N2B27 medium supplemented with 3 µM of GSK-3 inhibitor was added to each well. 150 µl of medium was then replaced every 24h.

### Generation of mutant ES cells

Wild-type mES cells (EmbryoMax 129/SVEV) were used to generate different mutant lines following the CRISPR/Cas9 genome editing protocol described in (49). sgRNA targeting guides (Table S1) were cloned into a Cas9-T2A-Puromycin expressing plasmid containing the U6-gRNA scaffold (gift of A. Németh; Addgene plasmid, 101039). ES cells were transfected with 8 µg of plasmid using the Promega FuGENE 6 transfection kit and dissociated 48 h later for puromycin selection (1.5 µg ml^−1^). Clone picking was conducted 5–6 days later, and positive ES cell clones were assessed by PCR screen using the MyTaq PCR mix kit (Meridian Bioscience) and specific primers surrounding the targeted region (Table S2). Mutations were verified by Sanger sequencing (Table S3). For the *HoxB^Del(i9-13):Ins(CBS5-10)^*the inserted CTCF cassette was synthesized (GeneArt, ThermoFisher). The sequence of the cassette was defined by selecting 500bp around each of the six distinct CBS located in between *Hoxb9* and *Hoxb13*. In order to increase CRISPR guide efficiency, one base pair was added 10bp after and before CBS9.

### WISH of Gastruloids

Gastruloids were collected at the indicated stage and processed following a previously reported WISH procedure (30). They were fixed overnight in 4% PFA at 4°C and stored in methanol at −20°C until ready for processing. Each sample was rehydrated and prepared with Proteinase K (EuroBio) at 2.5 µg/ml for 2 min. They were then incubated in a blocking solution at 68°C for 4 h before incubation overnight with specific digoxigenin-labeled probes (Table S5) at a final concentration of 100-200 ng/ml. The next day, samples were washed and incubated with an anti-DIG antibody coupled to alkaline phosphatase (Roche, 1:3,000). Staining was performed with BM-Purple (Roche).

### Generation of *in silico* mutant files

A fasta file containing the sequence of the chromosome 11 including HoxBDel(i9-13):Ins(CBS5-10) and one with the sequence of the chromosome 11 including *HoxB^Del^* (renamed chr11_delB) were generated using the R package seqinr (50) with the sequence of chromosome 11 from mm10 from UCSC as a template. Both mutant chromosome 11 sequences available at https://doi.org/10.5281/zenodo.12723266 were concatenated with the sequences of all other autosomes, chr X, chr Y and mitochondrial DNA from mm10 (UCSC).

### CTCF and RAD21 ChIPmentation

ChIPmentation of CTCF and RAD21 was performed as described in (18). Fastqs of the RAD21 ChIPmentation of 120h wild-type gastruloids from (18) were retrieved from SRA (SRR19601466) and processed with datasets generated in this study. For data analysis, adapter and bad quality bases were removed from fastq files using cutadapt version 4.1 (51) (-a CTGTCTCTTATACACATCTCCGAGCCCACGAGAC -A CTGTCTCTTATACACATCTGACGCTGCCGACGA -q 30 -m 15). Filtered reads were aligned on the *in silico* mutant genome using bowtie2 version 2.4.5 with default parameters (51). No filtering for mutlimap reads was performed, for sequences that are duplicated between both alleles of chromosome 11. Coverage was computed with macs2 version 2.2.7.1 (52) (--call-summits --format BAMPE -B) removing PCR duplicates and then normalized to the million fragments after filtering used by macs2.

### CUT&RUN

Each ESC per genotype were washed into fresh PBS before dissociation. The dissociation was performed in 0.5mL StemPro Accutase (Gibco A1110501), 5min at 37°C. Cells were resuspended into 2mL of ESC media and span at 300g for 2min. Then, 0.5x10^6^ cells were processed according to the CUT&RUN protocol (53). First, cells were resuspended into 1mL of Wash Buffer (WB: 20mM HEPES-KOH pH7.5, 150 mM NaCl, 0.5mM Spermidin (Sigma S2626)) and bound to Concanavalin A-coated beads (BioMagPlus 86057) into Digitonin Wash Buffer (DWB) using a final concentration of 0.02% digitonin (Apollo APOBID3301) into WB. Second, cells were incubated with 0.5µg/100μl of anti-CTCF antibody (Active Motif 61311) in DWB at 4°C for 2hr. The pA-MNase was produce by the Protein Production and Structure Core Facility at EPFL and added at 0.5μl/100μl in Digitonin Wash Buffer for 1hr at 4°C. Third, cells were digested in Low Calcium Buffer (100mM CaCl2 into DWB). Reaction was stopped using cold 2X STOP Buffer (340mM NaCl, 4mM EGTA, 20mM EDTA, 0.02% Digitonin, 50µg RNase A (10mg/mL), 50µG Glycogen (5mg/mL)) for 5min. Finally, targeted chromatin was released for 30 min at 37 °C, then precipitated into 70% EtOH and stored at -20°C until sequencing libraries generation. Sequencing libraries were prepared with KAPA HyperPrep reagents (07962347001) with 2.5µl of adapters at 0.3µM and ligated for 1 h at 20 °C. Then, DNA was cleaned and size selected using 1:1 ratio of DNA:Ampure SPRI beads (A63881) followed by an additional 1:1 wash and size selection with HXB. The DNA was amplified for 16 cycles. Post-amplified DNA was cleaned and size selected using 1:1 ratio followed by an additional 1:1 wash and size selection with HXB. HXB is equal parts 40% PEG8000 (Fisher FIBBP233) and 5 M NaCl. Libraries were sequenced on NextSeq 500 or NovaSeq 6000. For data analysis, adapter and bad quality bases were removed from fastq files using cutadapt version 4.1 (51) (-a GATCGGAAGAGCACACGTCTGAACTCCAGTCAC -A GATCGGAAGAGCGTCGTGTAGGGAAAGAGTGT-q 30 -m 15). Filtered reads were aligned on the *in silico* mutant genome using bowtie2 version 2.4.5 (54) with adapted parameters (--very-sensitive --no-unal --no-mixed --no-discordant --dovetail -X 1000). No filtering for mutlimap reads was performed as sequences are duplicated between both alleles of chromosome 11. PCR duplicates were removed with Picard MarkDuplicates version 2.27.4 (55). Filtered BAM file was converted to BED with BEDTools version 2.30.0 (56). Coverage was computed with macs2 version 2.2.7.1 (52) (--nomodel --keep-dup all --shift -100 -- extsize 200 --call-summits -B) and then normalized to the million reads in peaks.

### HiChIP

The dataset was produced and extracted from (18), GSM6226299.

### RNA-seq

Murine E9.5 and E10.5 mutant or wild-type littermate embryos were dissected to remove the head. RNA was extracted using RNeasy plus micro kit (Qiagen) following the manufacturer recommendation. The E9.5 samples libraries are poly(A)-enriched and E10.5 are ribo-depleted. Both were prepared with stranded RNA TruSeq kits (Illumina) and sequenced 100bp single-read on a HiSeq 4000. 120h and 144h AA gastruloids were collected and RNA was extracted as in (30). Libraries were prepared with ISML kit from Illumina, except for two replicates of *HoxB^Del/Del(i9-13):Ins(CBS5-10):Inv(CBS5-10)^*clone 1 at 120h, which were prepared with the TruSeq Stranded mRNA kit from Illumina. Libraries were sequenced on a NextSeq 500 or NovaSeq 6000 or HiSeq 4000, paired-end 2 times 75bp. Fastqs of the two replicates of 120h and 144h wild-type gastruloids from (18) were retrieved from SRA (SRR19600485 and SRR19600486, SRR19600479 and SRR19600480) and processed with datasets generated in this study. For data analysis, adapter and bad quality bases were removed from fastq files using cutadapt version 1.18 (51) (-q 30 -m 15 and sequences for -a and -A were adapted to the library). Filtered reads were aligned on mm10 using STAR version 2.7.10a (57) with the ENCODE parameters and a custom gtf (https://doi.org/10.5281/zenodo.7510406). FPKM were computed with cufflinks version 2.2.1 (58, 59) using --max-bundle-length 10000000 --multi-read-correct --library-type "fr-firststrand" -b mm10.fa --no-effective-length-correction -M mm10_chrM.gtf. Counts from STAR and FPKM values were subsetted for protein coding genes and genes on chrM were excluded. For the embryo datasets, genes on chromosomes X and Y were also excluded. Differential expression analysis was performed with DESeq2 version 1.38.0 (60) on R version 4.2.2. PCA was computed using log2(1 + FPKM) using the 500 most variant genes. **scRNA-seq**. scRNA-seq of gastruloids at 144h were performed as in (61), with 10x Chemistry v3 as well as the analysis up to matrix generation. Then, analysis was performed using Seurat v4.3 (62) with R version 4.3.0. We first filtered out barcodes with less than 200 identified gene and genes identified in less than three cells. Low quality cells and potential doublets were removed by computing the mean UMI content and the percentage of mitochondrial genes and filtering out barcodes with less than 0.4 times the mean UMI or more than 2.5 times the mean UMI. Only barcodes between 0.05 and 8 percent of mitochondrial UMI were kept. Matrices were normalized and the cell cycle score (using the 2019 updated gene list from Seurat) from these filtered libraries were computed. Then we merged the different samples using the merge command by Seurat. The combined Seurat object was normalized, 2000 variable features were identified and the data was scaled and regressed by cell cycle score and percentage of mitochondrial reads. Principal components were then computed using variable genes falling within the 5th and 80th percentile of expression to limit batch effect as performed in (63). UMAP and k-nearest neighbors were computed using 30 principal components. Clustering was performed with a resolution 0.4 and cluster annotation was performed manually. The average expression of a gene in a cluster was computed as the sum of the raw counts from all cells divided by the sum of the total counts of all cells. This value was log transformed like with a pseudo count of 1 and a scale of 10^4^. The inferred expression distribution of *Hoxb13* in mutant neuronal cells was obtained with baredSC version 1.1.2 (32) (--xmax 3 --minNeff 200 --minScale 0.1 for 1 to 4 gaussians combined), for the joint inferred expression distribution of *Hoxb13* with *Hoxa9*, *Hoxb9*, *Hoxc9* or *Hoxd* genes, similar parameters were used (--xmax 3 --ymax 3 --minNeff 200 -- minScalex 0.1 --minScaley 0.1 for 1 to 4 gaussians combined). Convergence of each MCMC was manually inspected using corner plots and ACF.

### minION sequencing

nCATS was performed as in (64) with a pool of 4 guides and a pool of 8 guides (Table S1). Base calling was computed with Guppy version 5.0.16+b9fcd7b from Oxford Nanopore (--flowcell FLO-MIN106 --kit SQK-LSK109 --fast5_out). Mapping was performed on mm10 or on the *in silico* mutant genome using minimap2 version 2.28 (65) (-ax splice). Non primary alignments were removed with samtools version 1.16.1 (66) and coverage was generated by BEDTools version 2.30.0 (56).

### Gene distance analysis

Gene distances were extracted from Ensembl Release 111. For each species of the database, genes with external gene names matching *Hoxb1*, *Hoxb9*, *Hoxb13* or derivatives were extracted, as well as all genes registered as homologous to the three mouse genes. Only couple of genes which were on the same chromosome were considered and the difference between the coordinate of the 3’ end of the first exon was used.

### Sequence conservation

Sequence conservation between the mouse and other vertebrate *HoxB* loci was visualized using the Vertebrate Multiz Alignment & Conservation maf file provided by UCSC for mm10. The Repeat Mask was also provided by UCSC. The sequence conservation between *mus musculus* and other strains was visualized using the 21-way Enredo-Pecan-Ortheus (EPO) multiple alignments provided by Ensembl. This sequence conservation was computed on mm39. To evaluate distance conservation between the various mouse strains, gene annotations were downloaded from Ensembl version 102 (mm10) and centered on *Hoxb9*.

### Data analysis

All NGS analyses were computed using the facilities of the Scientific IT and Application Support Center of EPFL. The genomic tracks were displayed using pyGenomeTracks version 3.9 (67, 68). The quantifications were plotted with R version 4.4.0 (https://www.R-project.org/) and ggpubr version 0.6.0 (69).

### Code availability

All scripts are available at https://github.com/lldelisle/scriptsForLopezDelisleZakanyBochatonEtAl2024.

### Data availability

All raw and processed datasets are available in the Gene Expression Omnibus (GEO) repository under accession number GSE272483.

## SUPPLEMENTARY FIGURES

**Figure S1.**
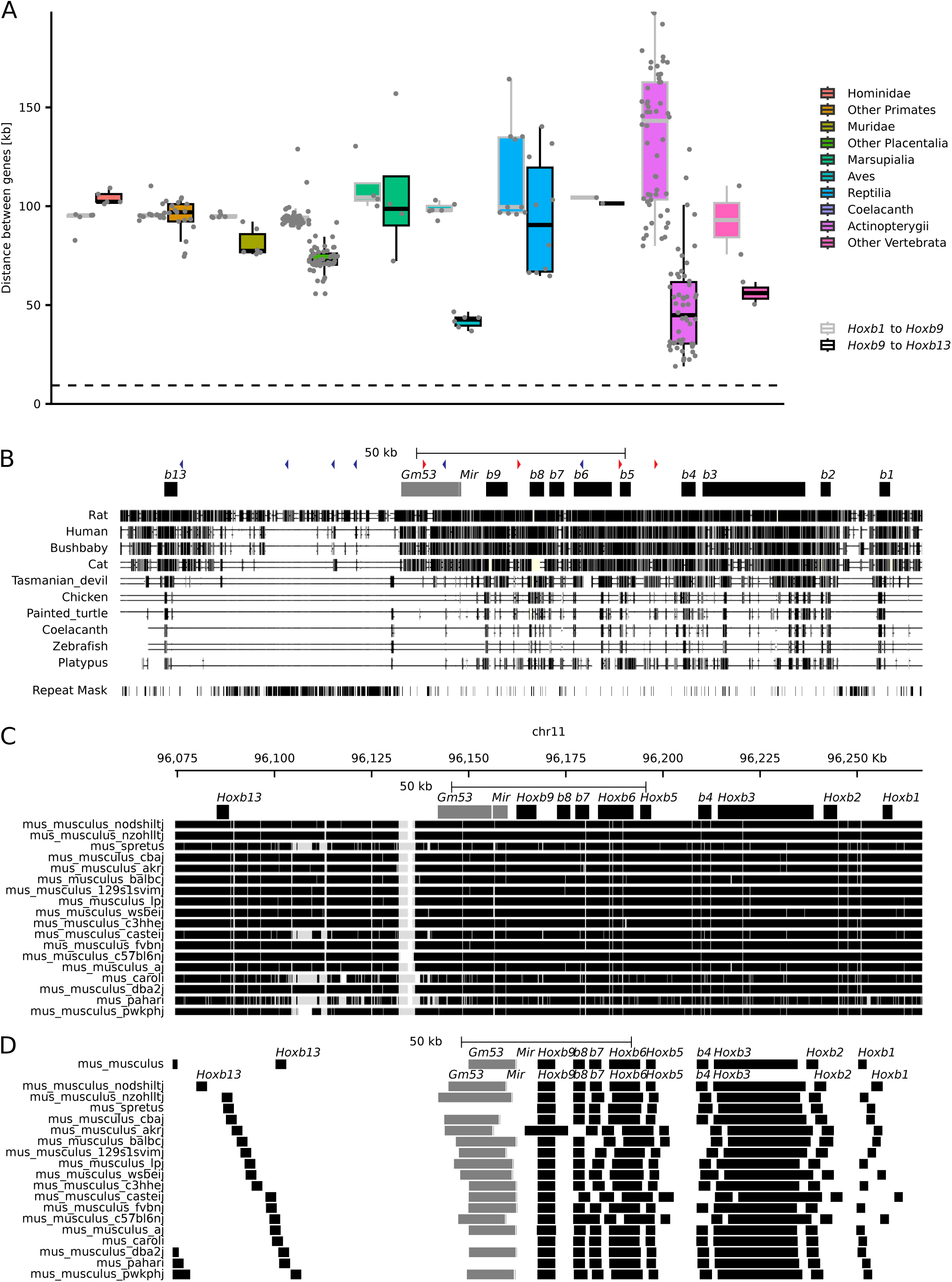
Conservation of the large DNA interval between *Hoxb9* and *Hoxb13*. (**A**) Boxplot showing the distance between either *Hoxb1* to *Hoxb9* (light grey) or *Hoxb9* to *Hoxb13* (black) for different species of Hominidae i.e., Bonobo, Chimpanzee, Gorilla, Human, Sumatran orangutan; ‘Other primates’ are Black snub-nosed monkey, Bolivian squirrel monkey, Bushbaby, Coquerel’s sifaka, Crab-eating macaque, Drill, Golden snub-nosed monkey, Greater bamboo lemur, Ma’s night monkey, Macaque, Mouse Lemur, Olive baboon, Panamanian white-faced capuchin, Pig-tailed macaque, Sooty mangabey, Vervet-AGM, White-tufted-ear marmoset; Muridae are Algerian mouse, Mouse, Rat, Ryukyu mouse, Shrew mouse, Steppe mouse ; Other Placentalia are American bison, American mink, Arabian camel, Arctic ground squirrel, Beluga whale, Blue whale, Cat, Chacoan peccary, Chinese hamster CHOK1GS, Cow, Degu, Dingo, Dog, Domestic yak, Donkey, Elephant, Eurasian red squirrel, Ferret, Giant panda, Goat, Golden Hamster, Greater horseshoe bat, Guinea Pig, Horse, Hybrid - Bos Indicus, Kangaroo rat, Leopard, Lesser Egyptian jerboa, Lion, Long-tailed chinchilla, Microbat, Naked mole-rat female, Narwhal, Northern American deer mouse, Pig, Pig - Bamei, Pig - Jinhua, Pig - Landrace, Pig - Pietrain, Pig - Tibetan, Pig USMARC, Prairie vole, Rabbit, Red fox, Sheep, Siberian musk deer, Sperm whale, Squirrel, Upper Galilee mountains blind mole rat, Vaquita, Wild yak, Yarkand deer; Marsupialia: Common wombat, Koala, Opossum, Tasmanian devil; Aves are: Chicken, Duck, Golden eagle, Japanese quail, Kakapo, Pink-footed goose; Reptilia are: Abingdon island giant tortoise, Argentine black and white tegu, Australian saltwater crocodile, Chinese softshell turtle, Eastern brown snake, Goodes thornscrub tortoise, Indian cobra, Mainland tiger snake, Painted turtle, Three-toed box turtle; Coelacanth; Actinopterygii are: Asian bonytongue, Atlantic cod, Atlantic herring, Barramundi perch, Bicolor damselfish, Brown trout, Burton’s mouthbrooder, Channel bull blenny, Channel catfish, Chinese medaka, Chinook salmon, Climbing perch, Clown anemonefish, Coho salmon, Common carp, Denticle herring, Eastern happy, Electric eel, Fugu, Gilthead seabream, Goldfish, Greater amberjack, Guppy, Huchen, Indian medaka, Japanese medaka HdrR, Javanese ricefish, Large yellow croaker, Lumpfish, Lyretail cichlid, Mangrove rivulus, Mexican tetra, Midas cichlid, Nile tilapia, Northern pike, Orange clownfish, Paramormyrops kingsleyae, Pike-perch, Pinecone soldierfish, Platyfish, Rainbow trout, Red-bellied piranha, Reedfish, Sheepshead minnow, Siamese fighting fish, Spiny chromis, Spotted gar, Tetraodon, Tiger tail seahorse, Turbot, Yellowtail amberjack, Zebra mbuna, Zebrafish, Zig-zag eel; Other Vertebrata are: Elephant shark, Platypus. The horizontal black dashed line represents the *Hoxb9* to *Hoxb13* distance in the *HoxB^Del(i9-13)^* allele for comparison. (**B**) Extent of sequence conservation between the mouse genome (mm10) and selected representative species: Rat (rn5), Human (hg19), Bushbaby (otoGar3), Cat (felCat5), Tasmanian devil (sarHar1), Chicken (galGal4), Painted turtle (chrPic1), Coelancanth (latCha1), Zebrafish (danRer7) and Platypus (ornAna1). Last track shows the repeat mask from UCSC showing interspersed repeats and low complexity DNA sequences. (**C**) Sequence conservation between the mouse reference genome (mm39) and 15 Mus musculus strains and Algerian mouse (Mus spretus), Ryukyu mouse (Mus caroli) and Shrew mouse (Mus pahari). The different genomes were ordered by decreasing *Hoxb13*-*Hoxb9* distance (see D). (**D**) Comparison of the *Hoxb13*-*Hoxb9* distances among the same strains and species as in **C**. All gene annotations were centred on *Hoxb9* for an easier comparison.

**Figure S2.**
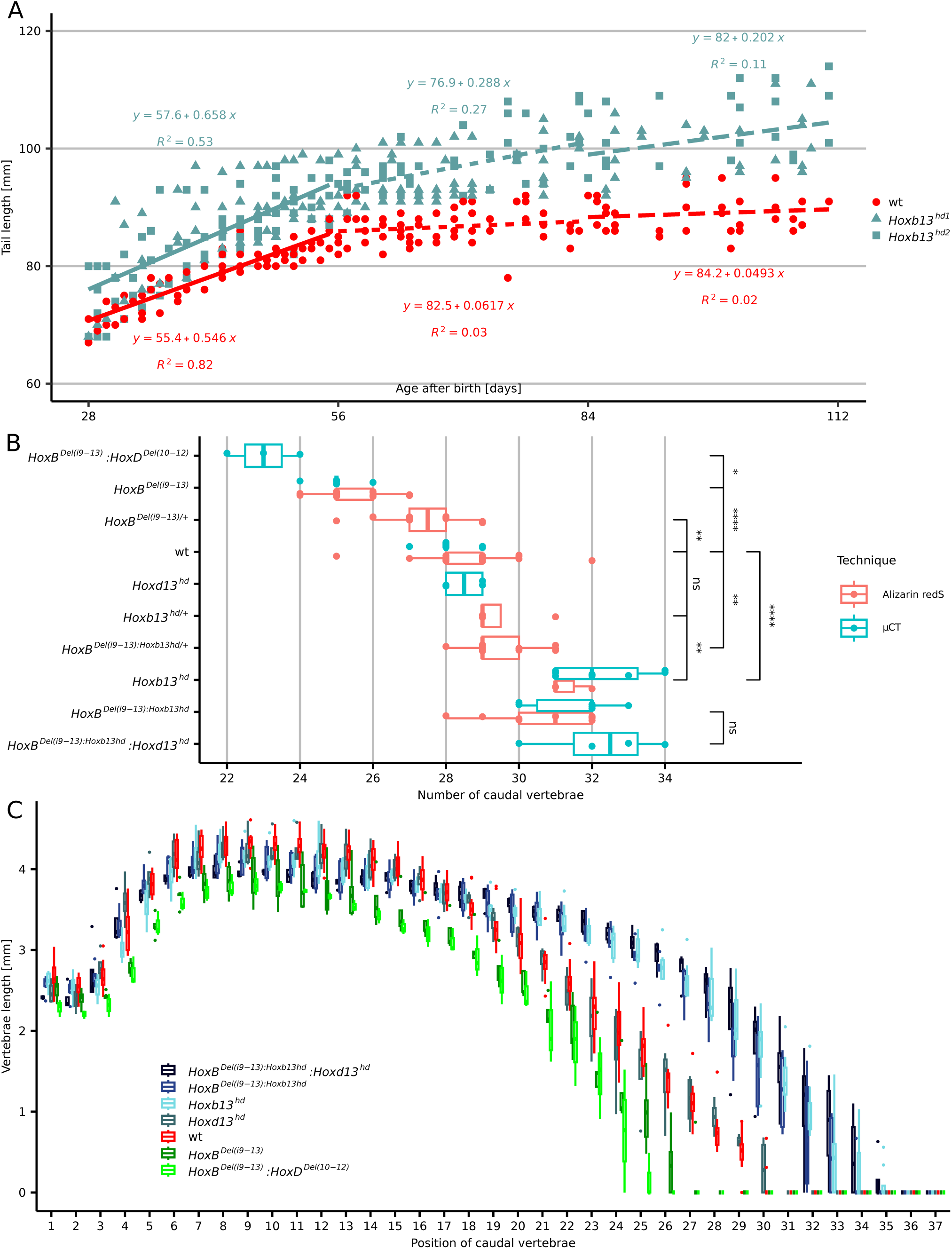
Quantifications of the variations in tail lengths amongst various alleles. (**A**) Tail growth trajectories of two different stocks of *Hoxb13^hd^* homozygous specimen (in blue, grouped together) and wild type siblings (in red, grouped together), during the second to fourth month of age. Linear regression equations were calculated on three equally-sized period of age, which are displayed above and below the regression lines. While the tail growth of wild-type animals stops at the end of the first period (linear coefficient below 0.1), the tail growth of mutant animals continues, leading to an increased difference of tail length along with the age. 175 measures of 15 wt animals, 138 measures of 9 *Hoxb13^hd1^*homozygous animals and 132 measures of 11 *Hoxb13^hd2^* homozygous animals were recorded. (**B**) Boxplots of caudal vertebra counts of ten genotypes (left) using either skeletal preparations (red) or µCT (blue). The numbers of caudal vertebrae are on the x axis. Brackets on the right indicate p-values obtained by pair-wise comparisons by Welch’s t-tests, (non-)significant differences are indicated by **** <0.0001, ***<0.001, **<0.01,*<0.05 or ns>0.05. In four cases both alizarin preparations and µCT analyses were available and the tightly similar distribution of counts between the different techniques validated the reliability of the observations where only the µCT data were obtained. Number of animals per genotype for Alizarin redS and µCT: *HoxB^Del(i9-13)^:HoxD^Del^*(*^10–12^*): 0 and 4; *HoxB^Del(i9-13)^* homozygous: 17 and 5; *HoxB^Del(i9-13)^* heterozygous: 14 and 0; wt: 28 and 12; *Hoxd13^hd^* homozygous: 0 and 4; *Hoxb13^hd^* heterozygous: 4 and 0; *HoxB^Del(i9-13):Hoxb13hd^* heterozygous: 14 and 0; *Hoxb13^hd^* homozygous: 3 and 8; *HoxB^Del(i9-13):Hoxb13hd^* homozygous: 11 and 6; *HoxB^Del(i9-13):Hoxb13hd^:Hoxd13^hd^* homozygous: 0 and 4. (**C**) Boxplot displaying the distribution of vertebrae length for each genotype at each position along the caudal region for animals analysed by µCT. When the caudal vertebrae did not exist, the size was set to 0. Six genotypes are plotted *HoxB^Del(i9-13):Hoxb13hd^:Hoxd13^hd^*, *HoxB^Del(i9-13):Hoxb13hd^*, *Hoxb13^hd^, Hoxd13^hd,^ wt, HoxB^Del(i9-13)^* and *HoxB^Del(i9-13)^*:HoxD*^Del^*(*^10–12^*).

**Figure S3.**
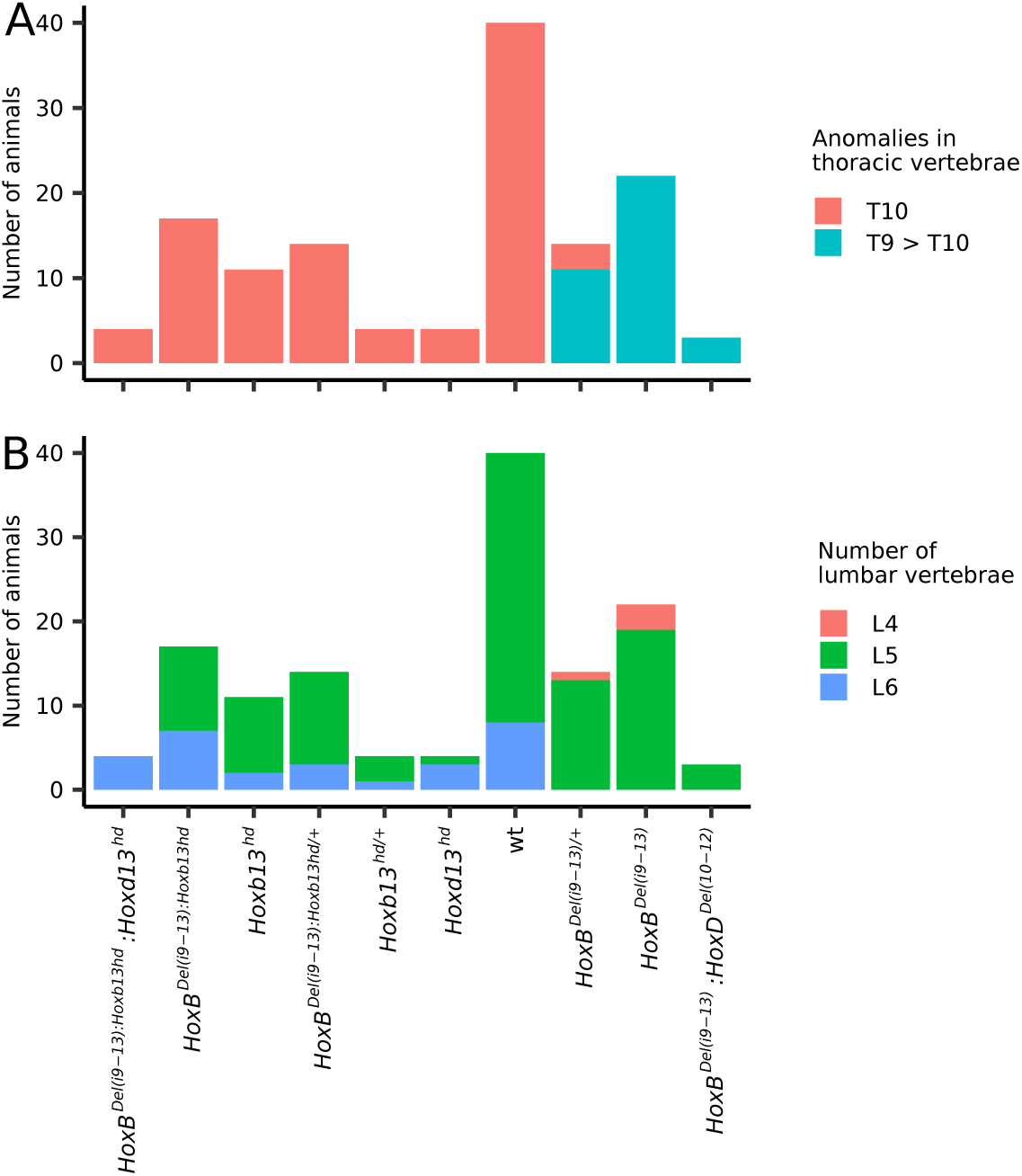
Quantification of vertebral anomalies. (**A**). Barplot showing the number of animals carrying either a classical ‘thoracic 10’ (T10) vertebral formula, or a T9 with an anticlinal morphology, which is usually typical of T10 (T9 > T10) for each of the 10 genotypes shown below. (**B**) Barplot showing the number of animals with the different numbers of lumbar vertebrae. The specimens are those reported in Figure S2B.

**Figure S4.**
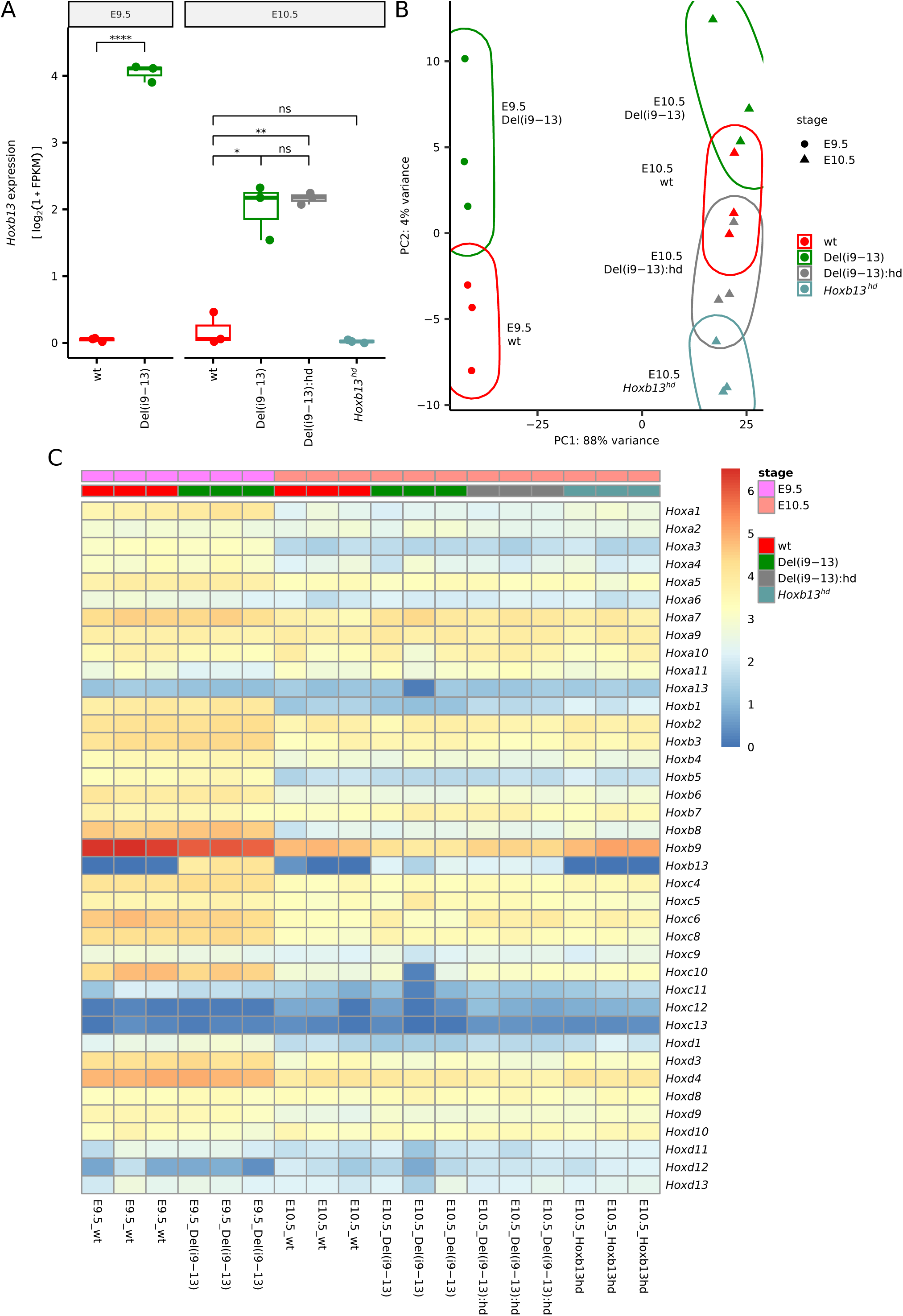
Quantification of *Hoxb13* transcripts. (**A**). Quantification of *Hoxb13* expression by RNA-seq in E9.5 and E10.5 headless embryos of the following genotypes (n=3): *HoxB^+/+^*(wt), *HoxB^Del(i9-13)^* homozygous (Del(i9-13)), *HoxB^Del(i9-13):Hoxb13hd^*homozygous (Del(i9- 13):hd) and *Hoxb13^hd^*homozygous (*Hoxb13^hd^*). The significance was assessed by the Wald test on counts with DESeq2, adjusted p-values: ns: p > 0.05, *: p <= 0.05, **: p <= 0.01, ***: p <= 0.001. (**B**) Projection on the two first principal components of all mouse samples analyzed by RNA-seq (same genotype as in **A**), at the same two time points. (**C**) Heatmap showing the expression of all *Hox* genes in the same samples as in A and B at the same two time points (below) (expressed in log2(1 + FPKM)).

**Figure S5:**
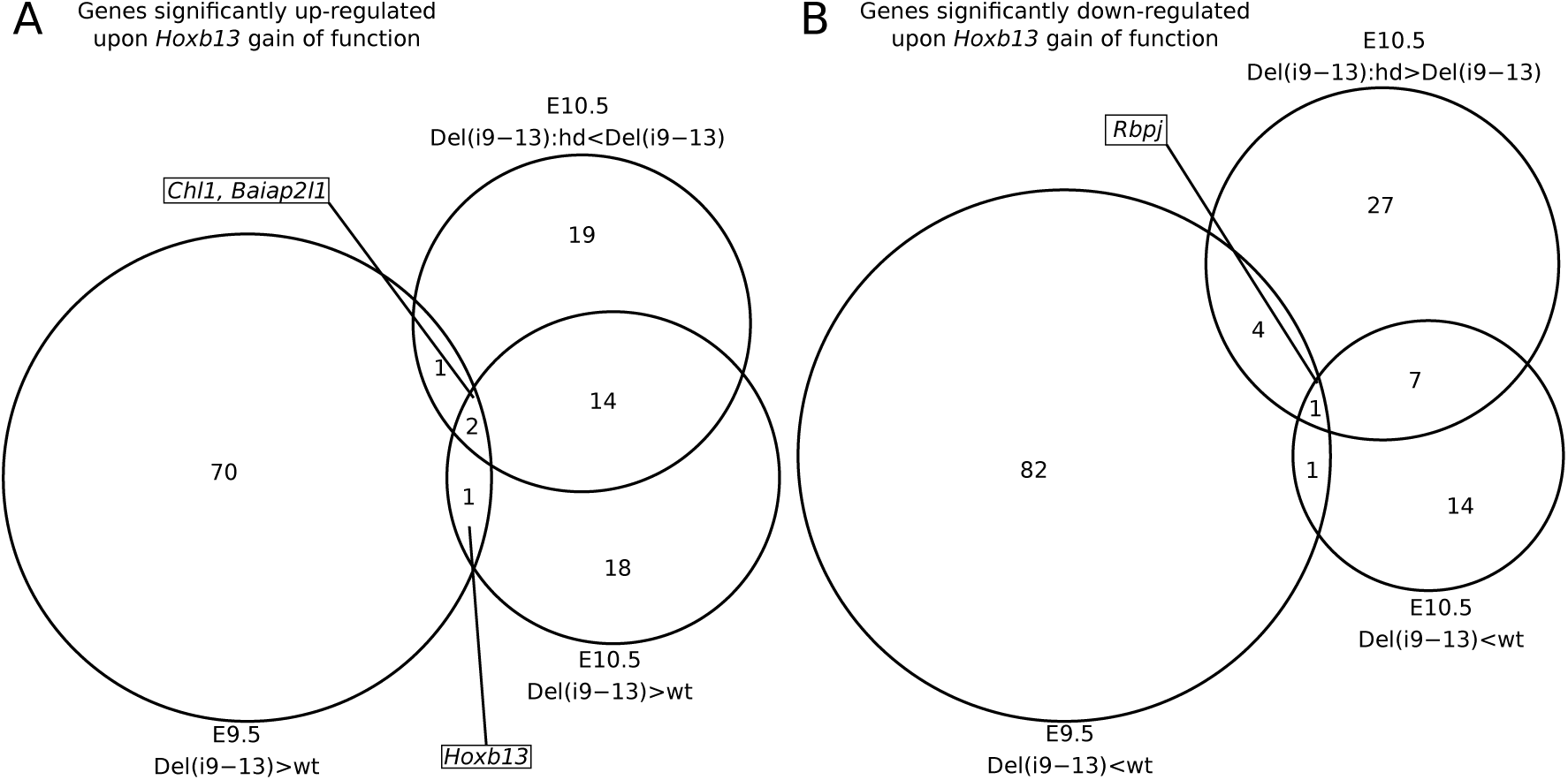
Global transcriptional changes associated with *Hoxb13* gain of function. (**A, B**) Euler diagrams highlighting the number of protein coding genes significantly upregulated (**A**) or downregulated (**B**) in gastruloids showing *Hoxb13* gain of function. The different datasets used form comparisons are: E9.5, wt *versus* Del(i9-13); E10.5, wt *versus* Del(i9-13); E10.5 Del(i9-13)hd *versus* Del(i9-13). Only two genes were significantly upregulated with *Hoxb13* gain of function (*Chl1*, *Baiap2l1*). A single gene is significantly downregulated with *Hoxb13* gain of function: *Rbpj.* No gene was found overlapping between the two Euler diagrams. Genes were considered significantly expressed when adjusted p-value for two-sided Wald test < 0.05 and fold-change above 1.5 or below 0.67.

**Figure S6.**
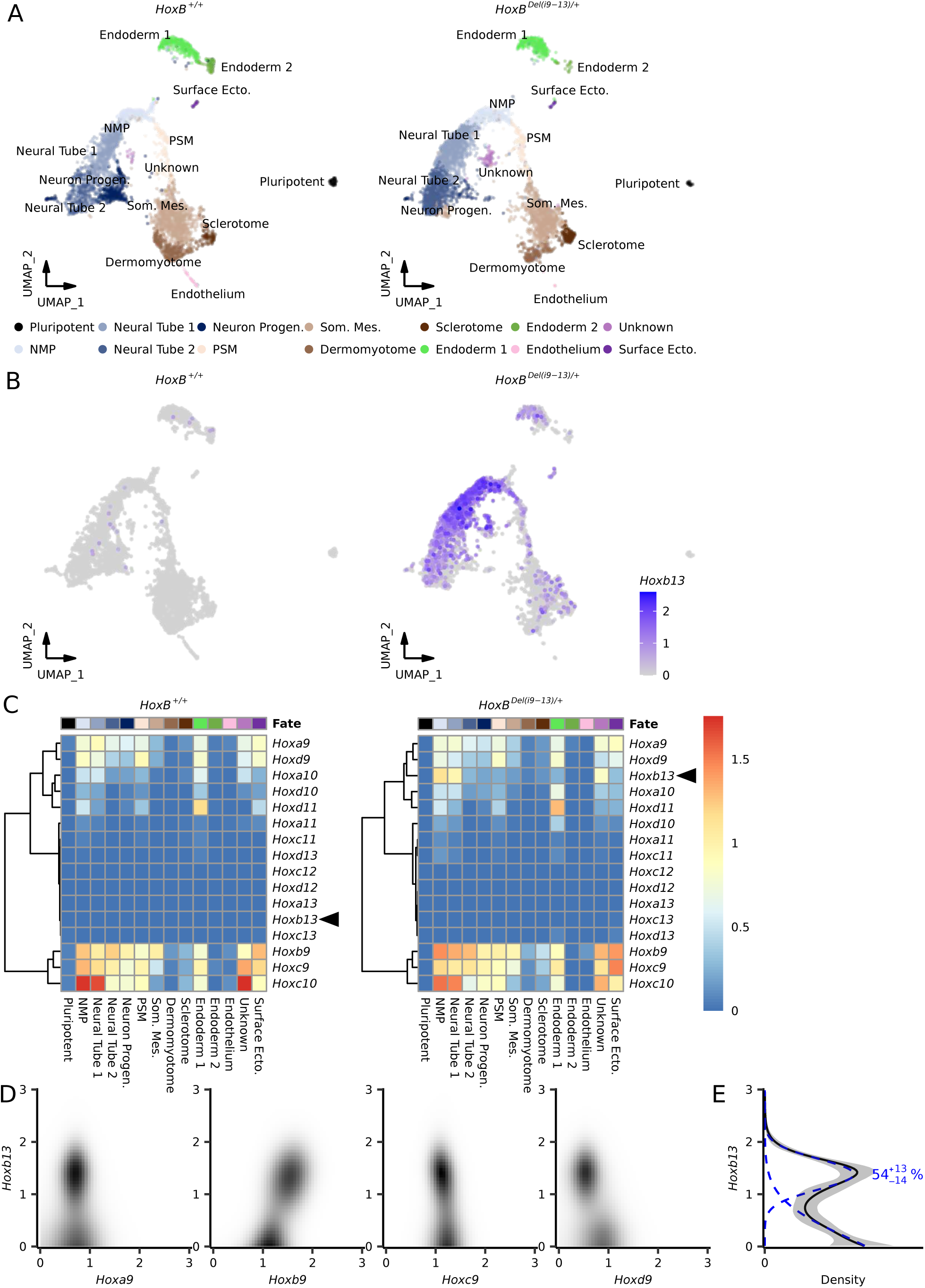
scRNA-seq analysis of gastruloids with deletion of the *HoxB* DNA spacer. (**A**) UMAP projections of single cells of wild-type (left) and *HoxB^Del(i9-13)/+^* (right) 144h gastruloids. Cells are coloured by clusters associated to cell types. (**B**) UMAP projection of single cells of wild-type (left) and *HoxB^Del(i9-13)/+^* (right) 144h gastruloids. Cells are coloured according to their levels of expression of *Hoxb13*. (**C**) Heatmap reporting the average expression of posterior *Hox* genes (group 9 to 13) on each cellular cluster either on wild-type (left) or in HoxB^Del(i9-13)/+^ (right) specimens. Hierarchical clustering was performed by using these related genes to highlight the strong similarity between all genotypes, except for *Hoxb13* (black arrowheads). (**D**) Heatmaps representing the inferred distribution of cells of the NMP cluster in *HoxB^Del(i9-13)/+^* gastruloids for the different levels of expression of either *Hoxb13* on the y-axis, or for *Hoxa9*, *Hoxb9*, *Hoxc9* and *Hoxd9* on the x-axis. The colour scale is log transformed in order to see a greater range of frequencies. Black bins represent a high proportion of cells whereas white bins indicate an absence of cells. (**E**) Inferred distribution of *Hoxb13* expression using a two-Gaussian model in the same cells as in (**D**). The average inferred distribution is in black and the 68 percentile confidence interval is in grey. In dashed blue lines are plotted the truncated Gaussians with the median characteristics (average and scale). The percentage indicates the proportion of cells in the Gaussian with the highest expression.

**Figure S7:**
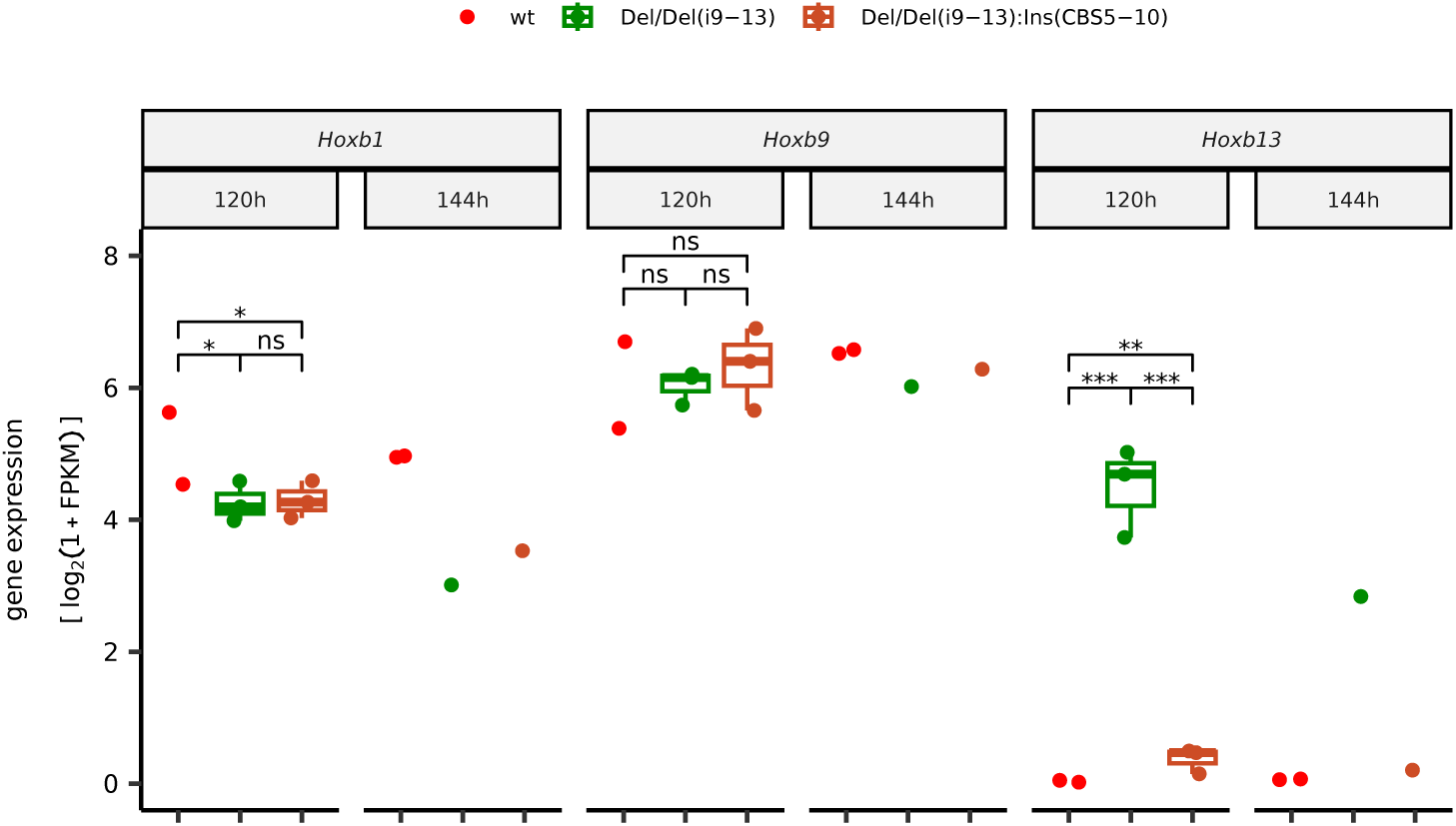
Quantification of *Hoxb* genes expression in gastruloids. Quantification of expression of *Hoxb1* (left), *Hoxb9* (middle) and *Hoxb13* (right) in gastruloids at 120h and 144h, in *HoxB^+/+^* (wt from (18)), *HoxB^Del/Del(i9-13)^* (Del/Del(i9-13)) and *HoxB^Del/Del(i9-13):Ins(CBS5-10)^* (Del/Del(i9-13):Ins(CBS5-10)). The significance was assessed by the Wald test on counts with DESeq2 at 120h only. The p-values were not corrected for multiple tests (ns: p > 0.05, *: p <= 0.05, **: p <= 0.01, ***: p <= 0.001).

**Figure S8:**
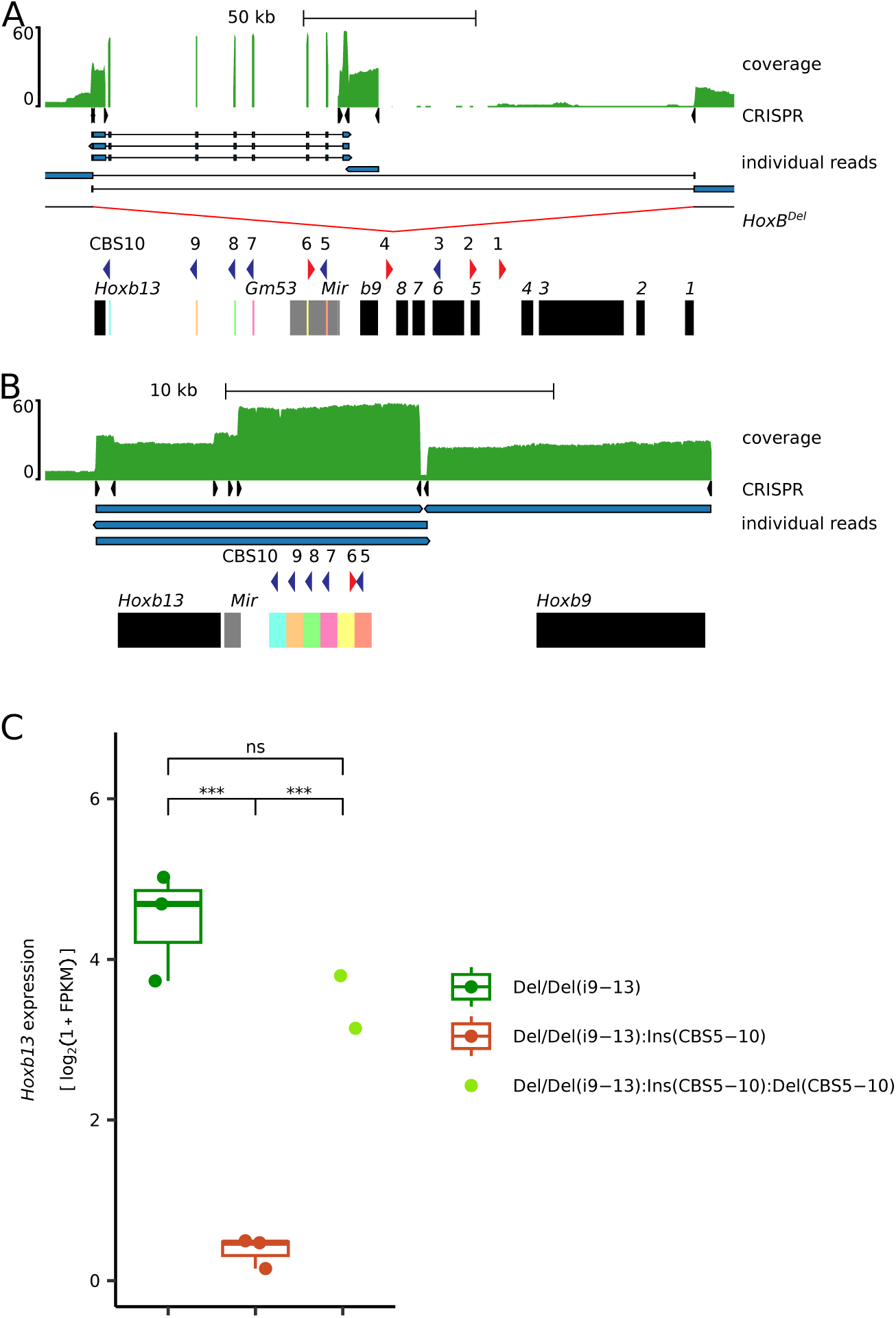
Controls for the Del(i9-13):Ins(CBS5-10) allele. (**A-B**) Control of the Del(i9-13):Ins(CBS5-10) allele by nCATS. First track shows the coverage obtained by nCATS when mapped on mm10 (**A**) or on the reconstructed mutant genome (**B**). Second track shows the CRISPR guides which were used in the nCATS experiment. Most reads are between two guides. Third track shows the mapping of individual reads selected because they cover the cassette or the *Hoxb9* gene or support the deletion of the *HoxB* cluster on the other allele (last two reads in **A**). Each coloured block containing a CBS corresponds to the flanking region in the wild-type allele (see **A**). (**C**) Quantification of *Hoxb13* RNAs in gastruloids at 120h in *HoxB^Del/Del(i9-13)^* (Del/Del(i9-13), n = 3), *HoxB^Del/Del(i9-13):Ins(CBS5-10)^* (Del/Del(i9-13):Ins(CBS5-10), n = 3) and *HoxB^Del/Del(i9-13):Ins(CBS5-10):Del(CBS5-10)^* (Del/Del(i9-13):Ins(CBS5-10):Del(CBS5-10), n = 2). The significance was assessed by the Wald test on counts with DESeq2. The p-values were not corrected for multiple tests (ns: p > 0.05, *: p <= 0.05, **: p <= 0.01, ***: p <= 0.001). The level of *Hoxb13* expression is restored when the CBS are deleted.

**Figure S9:**
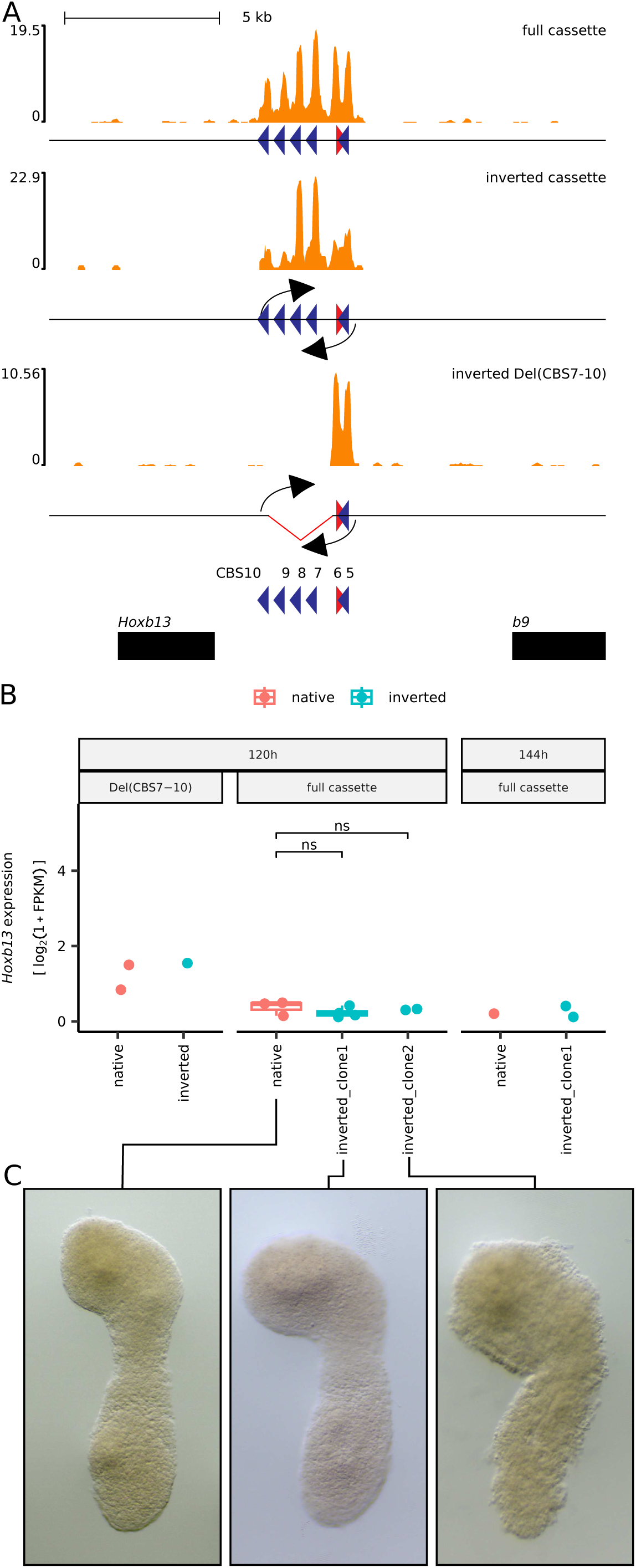
Inversion of CTCF sites in the inserted cassette. (**A**) CTCF profiles by CUT&RUN using ES cells on the *in silico* reconstructed genome of the following genotypes: *HoxB^Del/Del(i9-13):Ins(CBS5-10)^* (full cassette inserted, top track), *HoxB^Del/Del(i9-13):Ins(CBS5-10):Inv(CBS5-10)^* (full cassette inserted and then inverted) and HoxB*^Del/Del(i9-13):Ins(CBS5-10):Inv(CBS5-10):Del(CBS7-10)^* (full cassette inserted, then inverted, then deleted for CBS7-10). The arrows indicate the inversion sites, yet the CBS are shown in their ‘native’ (pre-inversion) orientations. The positions of *Hoxb9* and *Hoxb13* is shown below (black rectangles). (**B**) Quantification of *Hoxb13* expression in log2(1 + FPKM) in gastruloids at 120h (left) and 144h (right) with the CTCF cassette inserted either in the native orientation (red) or in the inverted orientation (blue). Genotypes are from left to right: *HoxB^Del/Del(i9-13):Ins(CBS5-10):Del(CBS7-10)^* (n = 2), *HoxB^Del/Del(i9-13):Ins(CBS5-10):Inv(CBS5-10):Del(CBS7-10)^* (n = 1), *HoxB^Del/Del(i9-13):Ins(CBS5-10)^* (n = 3), *HoxB^Del/Del(i9-13):Ins(CBS5-10):Inv(CBS5-10)^* (clone 1, n = 4), *HoxB^Del/Del(i9-13):Ins(CBS5-10):Inv(CBS5-10)^* (clone 2, n = 2), *HoxB^Del/Del(i9-13):Ins(CBS5-10)^* (n = 1), *HoxB^Del/Del(i9-13):Ins(CBS5-10):Inv(CBS5-10)^* (n = 2). The significance at 120h for the inversion was assessed by the Wald test on counts with DESeq2. The p-values were not corrected for multiple tests (ns: p > 0.05). (**C**) Whole mount *in situ* hybridisation with a *Hoxb13* antisense probe of 120h gastruloids of *HoxB^Del/Del(i9-13):Ins(CBS5-10)^* (left) and two different clones of HoxB*^Del/Del(i9-13):Ins(CBS5-10):Inv(CBS5-10)^* (middle and right).

---

**Table S1:**
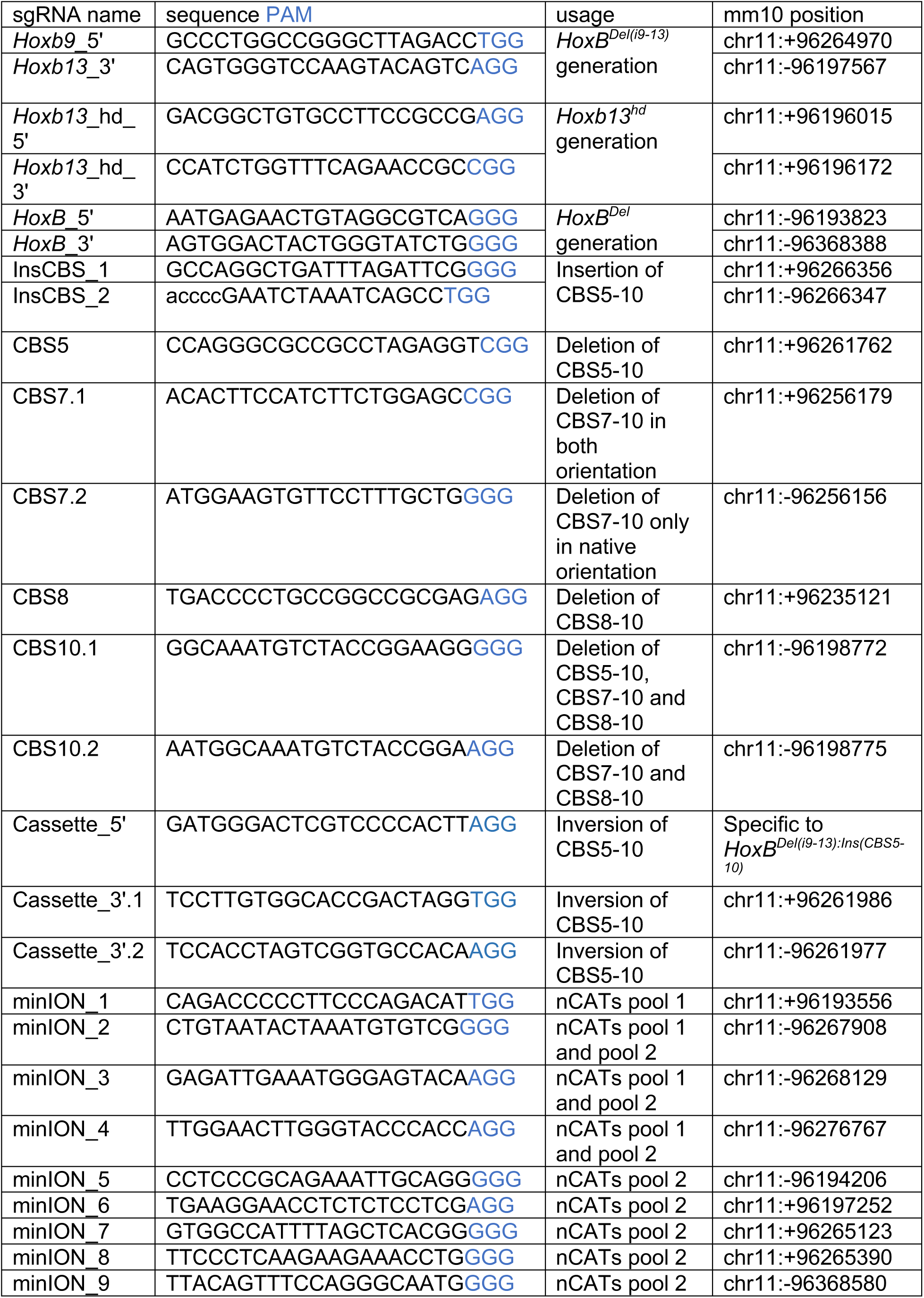
CRISPR sgRNAs.

**Table S2:**
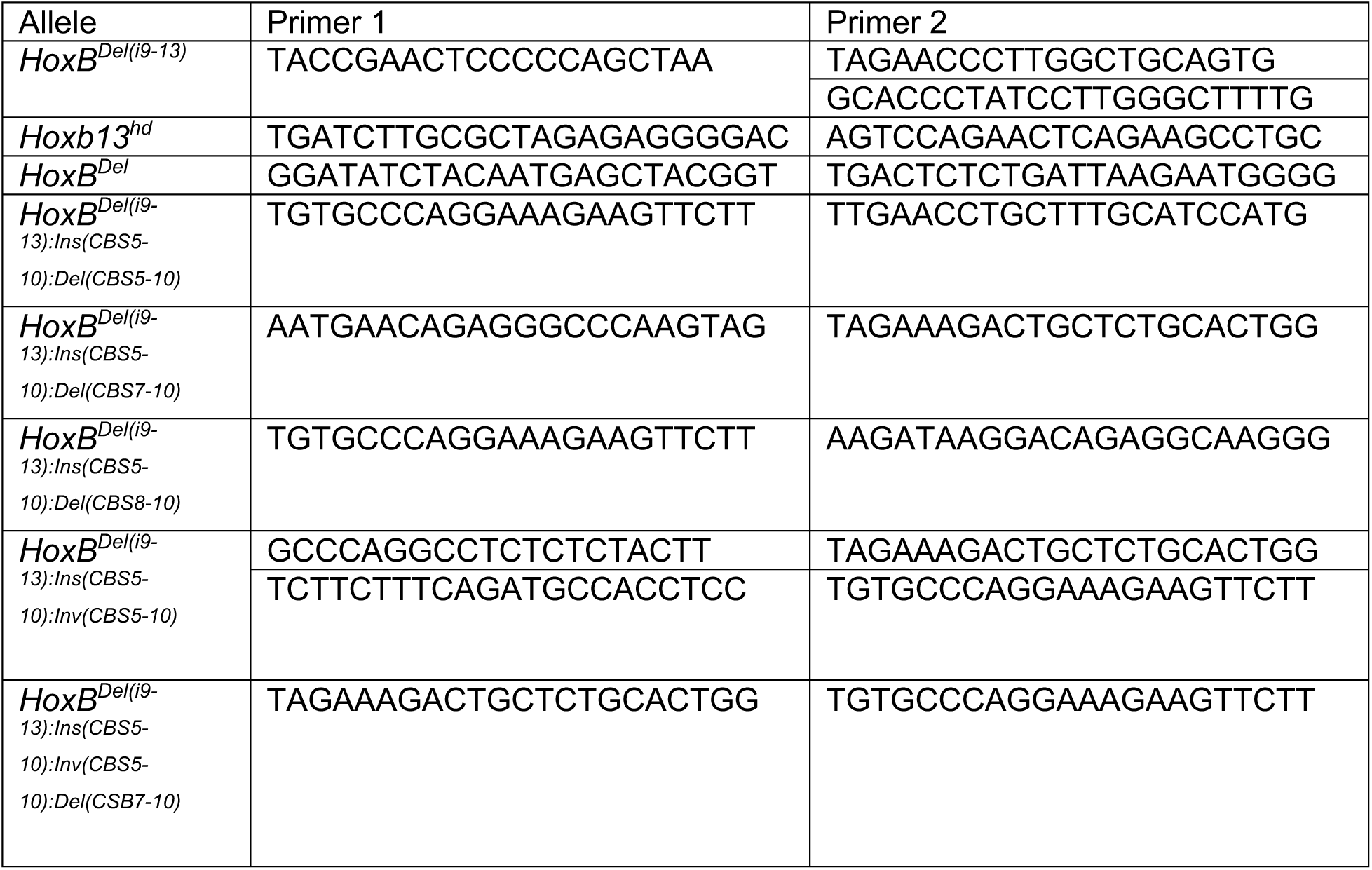
Genotyping primers.

**Table S3:**
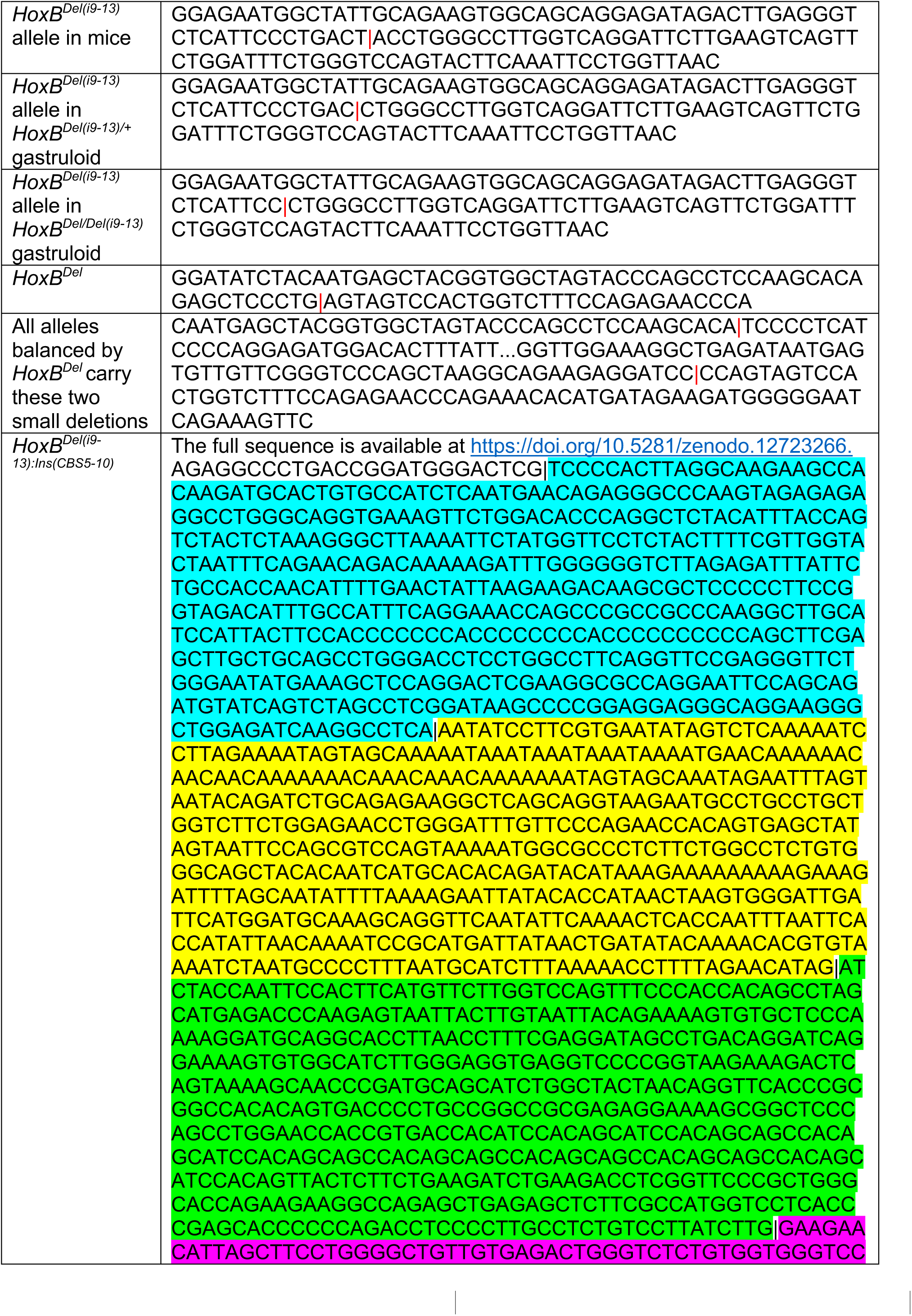
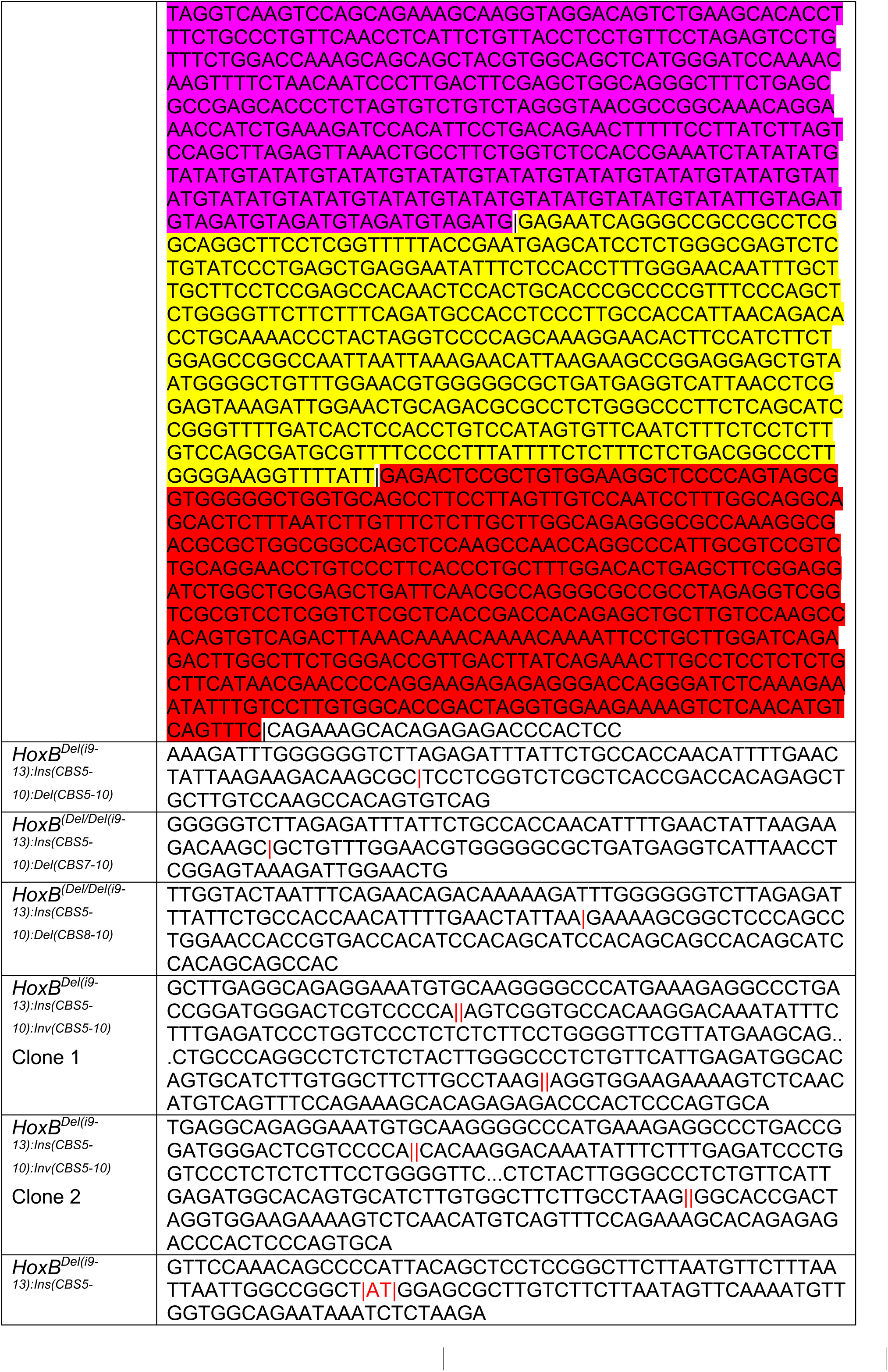
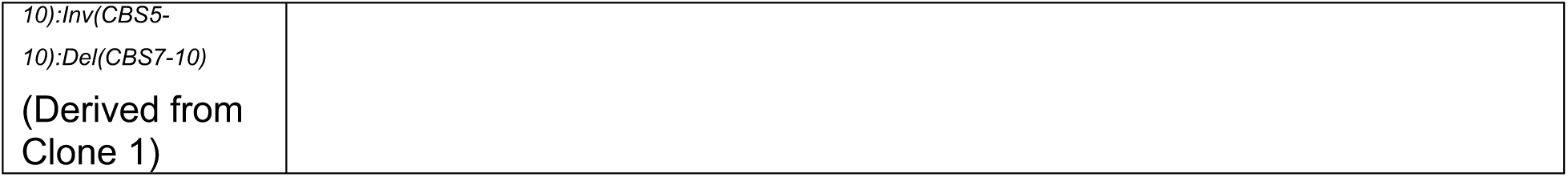
Sequences of various breakpoints.

**Table S4:**
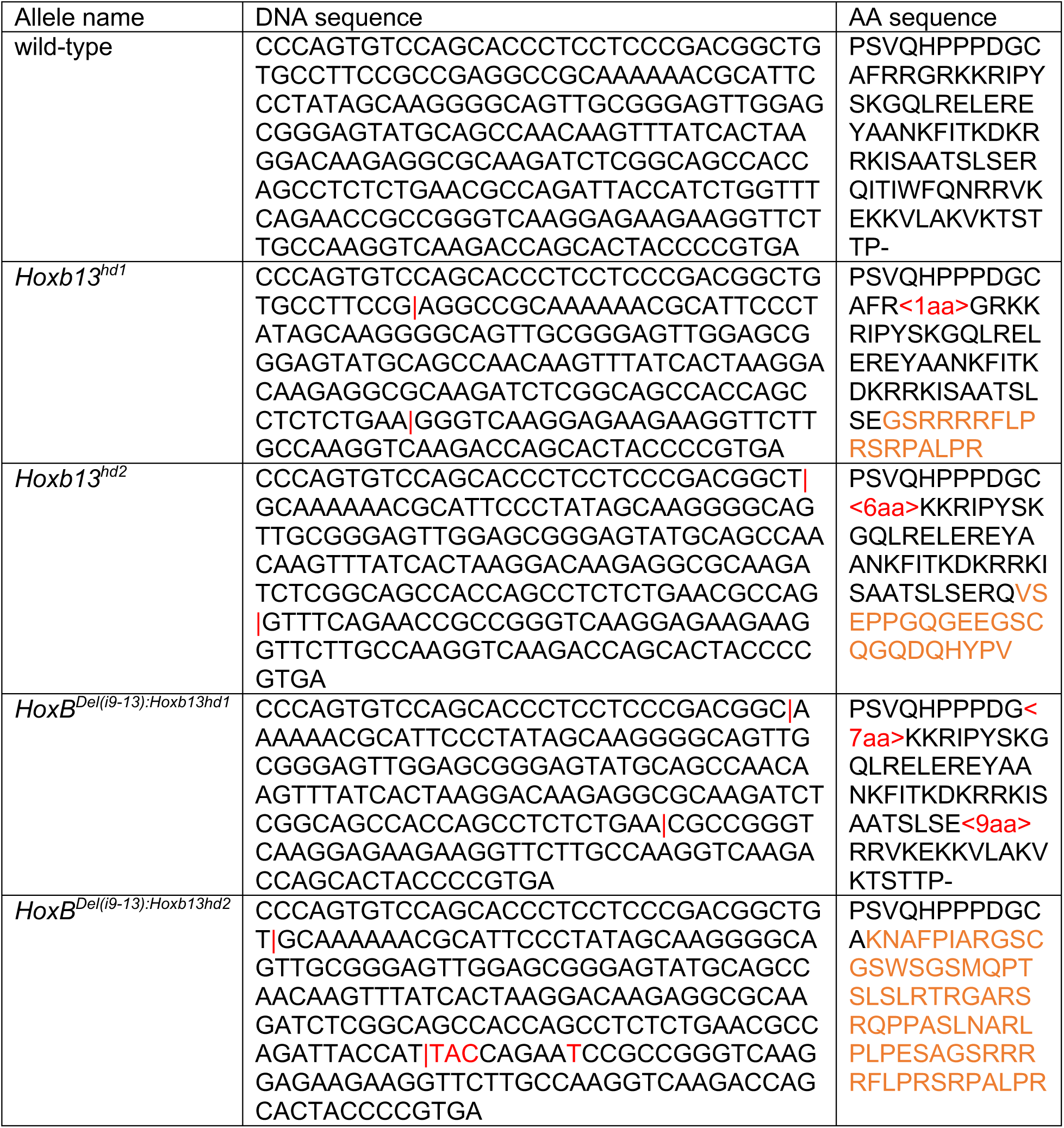
Homeobox/domain sequences of the two *Hoxb13^hd^* null alleles.

**Table S5:**
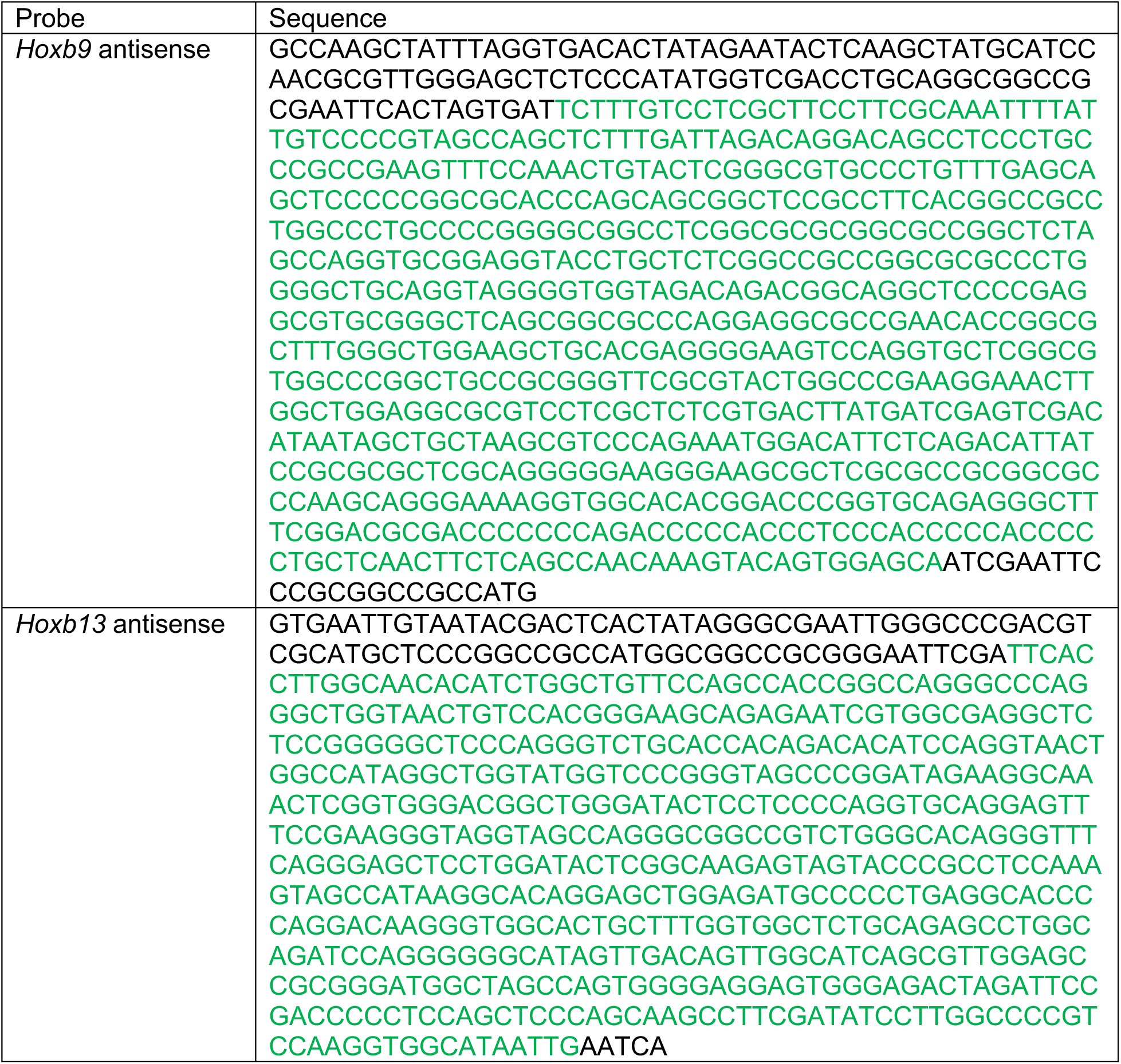
Probes for WISH.

